# A Novel Clustering Method for Patient Stratification

**DOI:** 10.1101/073189

**Authors:** Hongfu Liu, Rui Zhao, Hongsheng Fang, Feixiong Cheng, Yun Fu, Yang-Yu Liu

## Abstract

Patient stratification or disease subtyping is crucial for precision medicine and personalized treatment of complex diseases. The increasing availability of high-throughput molecular data provides a great opportunity for patient stratification. In particular, many clustering methods have been employed to tackle this problem in a purely data-driven manner. Yet, existing methods leveraging high-throughput molecular data often suffers from various limitations, e.g., noise, data heterogeneity, high dimensionality or poor interpretability. Here we introduced an Entropy-based Consensus Clustering (ECC) method that overcomes those limitations all together. Our ECC method employs an entropy-based utility function to fuse many basic partitions to a consensus one that agrees with the basic ones as much as possible. Maximizing the utility function in ECC has a much more meaningful interpretation than any other consensus clustering methods. Moreover, we exactly map the complex utility maximization problem to the classic *K*-means clustering problem with a modified distance function, which can then be efficiently solved with linear time and space complexity. Our ECC method can also naturally integrate multiple molecular data types measured from the same set of subjects, and easily handle missing values without any imputation. We applied ECC to both synthetic and real data, including 35 cancer gene expression benchmark datasets and 13 cancer types with four molecular data types from The Cancer Genome Atlas. We found that ECC shows superior performance against existing clustering methods. Our results clearly demonstrate the power of ECC in clinically relevant patient stratification.

## Introduction

High-throughput technologies, such as next-generation sequencing, have enabled us to rapidly accumulate a wealth of various molecular data types, including genome, transcriptome, proteome, and epigenome (*1*-*3*). Those massive genomics studies offer us great opportunities to characterize human pathologies and disease subtypes, identify driver genes and pathways, and nominate drug targets for precision medicine (*4*, *5*). In particular, development of novel computational approaches for patient stratification leveraging high-throughput molecular data would significantly facilitate precision medicine and personalized treatment, which target discrete molecular subclasses of complex diseases with specific genetic or epigenetic profiles (*4*).

*Clustering*, an unsupervised exploratory analysis, has been widely used for patient stratification or disease subtyping (*6*). However, traditional clustering algorithms, such as *K*-means, hierarchical clustering, and spectral clustering, suffer from noise, data heterogeneity and high dimensionality that are associated with high-throughput molecular data (*7*, *8*). *Ensemble clustering* (a.k.a. *consensus clustering*) can merge some individually generated basic partitions, and ensure the final consensus partition maximally agrees with the basic ones (*9*). This significantly helps us generate more robust clustering results, find bizarre clusters, better handle noise, outliers and sample variations, and integrate solutions from multiple distributed data sources (*9*). However, existing consensus clustering algorithms based on co-association matrix (*10*) are computationally expensive and require a large storage space, preventing them to handle high-throughput molecular data. Moreover, their interpretation of the consensus partition is often obscure.

## Results

### Methodology overview of ECC

Here we introduce a novel consensus clustering method, i.e. *Entropy-based Consensus Clustering* (ECC), for patient stratification. Consider an *n×m* matrix of molecular data of *n* subjects (or experiments, conditions, samples; corresponding to *n* rows) and *m* features (such as mRNAs; corresponding to *m* columns). Each subject can be represented by a point in the *m*-dimensional feature space, with different shapes representing different clusters the subjects belong to (**Fig. 1A**). There are three steps in the ECC pipeline. Step-1: We generate *r* basic partitions using *K*-means clustering with parameter *K* (i.e., the number of clusters) randomly chosen from 2 to 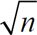 (see **Fig. 1B**) (*11*). Hereafter we call this kind of basic partitions generation strategy *Random Parameter Selection* (RPS). Note that in this step we can use any basic clustering method. Here we just choose *K*-means for its simplicity and high efficiency. In this work we choose *r* = 100 and find that larger *r* does not significantly improve the result. Step-2: We derive a binary matrix from each basic partition via 1-of*-K* coding, where *K* is the cluster number in this basic partition and only one element in each row is1, others are 0. We concatenate all those binary matrices into a large binary matrix (**Fig. 1C**). Step-3: We employ an entropy-based utility function to guide the fusion of all the *r* basic partitions into a consensus one (**Fig. 1D**). This is achieved by conducting *K*-means clustering on the binary matrix with a modified distance function and a user-defined cluster number *K*.

**Fig. 1.**
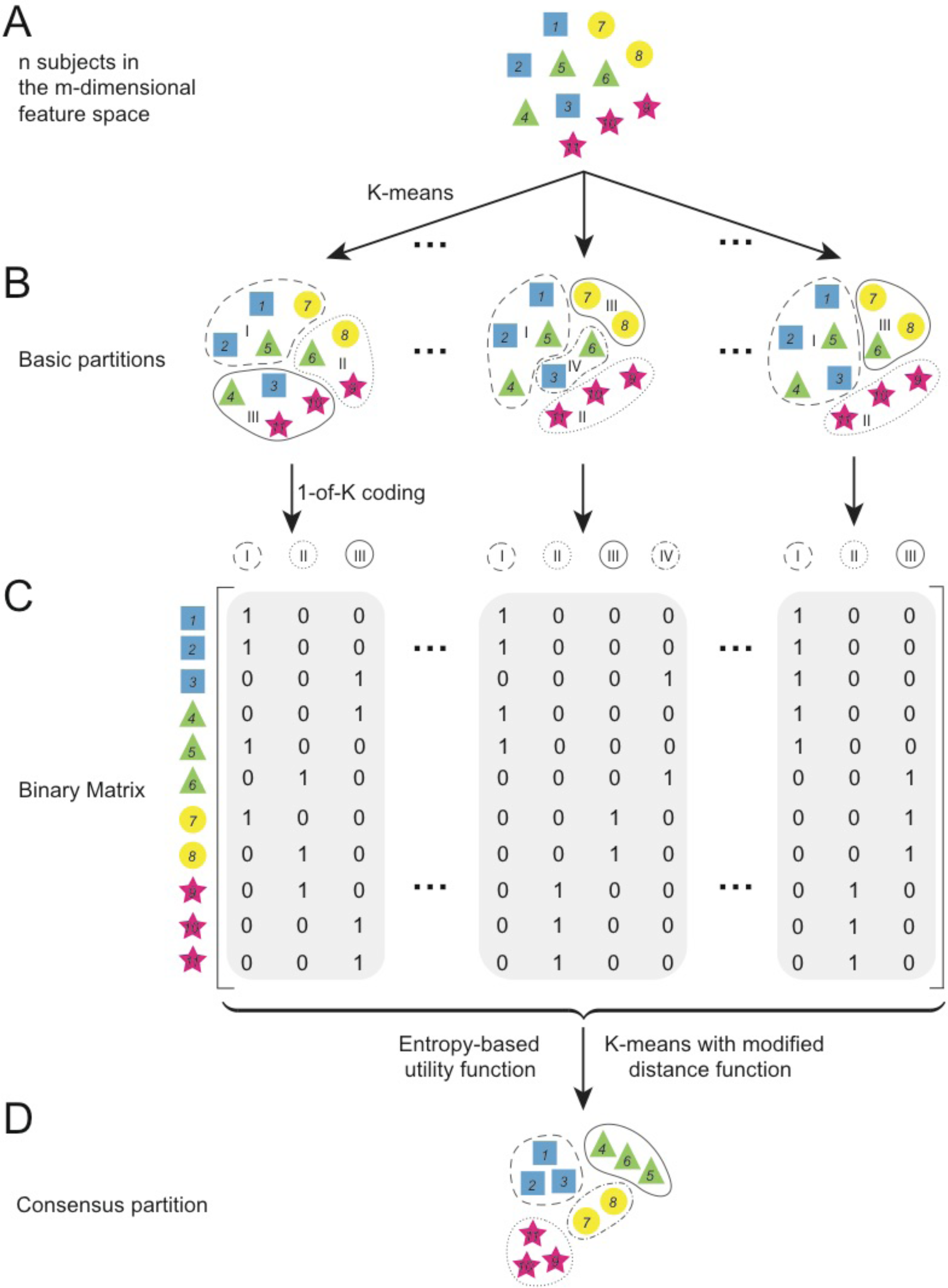
Schematic diagram of the ECC pipeline. (**A**) *n* subjects are presented by *n* points in the *m*-dimensional feature space. In this example, *n* = 11. The feature can be mRNA expression, Protein expression, or any other molecular data. Different shapes represent the subjects in different disease subtype (clusters). (**B**) *K*-means clustering is applied to the molecular data of the *n* subjects to obtain *r* basic partitions. For each basic partition, the cluster number *K* is randomly chosen from 2 to 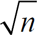, and we highlight the *K* clusters using dashed line, dotted line, solid line, etc. (**C**) Each basic partition is transformed into 1-of-*K* coding, where *K* is the cluster number in each basic partition and only one element in each row is 1, others are 0. Concatenating all the basic partitions in 1-of-*K* coding form yields a large binary matrix **B**, which is a new representation of the original molecular data. (**D**) A *K*-means clustering with modified distance function (derived from an entropy-based utility function) is conducted on the binary matrix **B** for the final consensus clustering. In this step, for synthetic and benchmark cancer gene expression datasets we set *K* to be the true cluster number. For the 13 TCGA cancer datasets, to fairly compare the ECC method and other clustering methods, we use the empirical number of clusters (subtypes) obtained from previous studies. For general molecular data when the empirical number of clusters is unknown, we can employ the cluster number estimation method in (*33*) to determine *K* for the final step of ECC.

Our ECC method has three key features. First, it solves the consensus clustering problem in a *utility* way, which has more meaningful interpretation than any other consensus clustering methods. Here the utility function is applied to quantify the similarity between each of the *r* basic partitions and the consensus one. Maximizing the utility function requires us to find a single consensus partition that agrees with the basic ones as much as possible. Second, we uncover a remarkable equivalence relationship between an entropy-based utility function and a *K*-means distance function so that the complex utility maximization problem can be efficiently solved by the classic *K*-means method with a modified distance function (see **Supplementary Materials Sec. I.A**). Consequently, both the time and space complexity of ECC are linear in *n* (see **Supplementary Materials Sec. I.B**). This dramatically improves the efficiency of ECC in real-world applications (*11*). Finally, ECC can naturally integrate multiple molecular data types measured from the same set of subjects, and easily handle missing values without any imputation (see **Materials and Methods**). This significantly increases the power of ECC in clinically relevant patient stratification.

To demonstrate that ECC indeed outperforms existing clustering methods, we compared the performance of ECC with five traditional clustering methods: Agglomerative Hierarchical Clustering with Average Linkage (AL), Single-Linkage (SL) and Complete-Linkage (CL), *K*-means Clustering (KM), and Spectral Clustering (SC); and two state-of-the-art consensus clustering methods: the Link-based Cluster Ensemble (LCE) and Approximate SimRank-based (ASRS) methods (*12*).

### Evaluation using synthetic data

We first applied all those clustering methods to synthetic gene expression datasets with built-in cluster structure generated through a well-established dynamical gene regulation model (see **Supplementary Materials Sec. II**) (*13*). For fair comparison, we used two external indices of clustering validity: *R_n_* (Normalized Rand Index) and *NMI* (Normalized Mutual Information) to objectively evaluate the performance of different clustering methods (*14*). Both *R_n_* and *NMI* are positive cluster validity indices that estimate the quality of clustering results with respect to the underlying cluster structure of the data (see **Materials and Methods**). We found that ECC generally outperforms other methods in terms of its robustness against noise (see **Supplementary Materials Sec. IV.A**).

### Evaluation using benchmark cancer gene expression data

We then evaluated ECC and other clustering methods on 35 widely usedbenchmark cancer gene expression datasets(*15*) (**Supplementary Materials Sec. III.A**). The detailed description of the 35 datasets was provided in **Supplementary Table S2**. **Fig. 2A** shows the clustering performance of different algorithms measured by *NMI*. We found that for most datasets, the three consensus clustering methods (LCE, ASRS, and ECC) are superior to the five traditional clustering methods. Moreover, our ECC method achieves promising results on several datasets by a large margin, such as dataset-5 (*Armstrong-2002-v2*), dataset-9 (*Yeoh-2001-v1*), dataset-10 (*Chowdary-2006*) and dataset-13 (*Golub-1999-v1*). Although LCE and ASRS yield reasonable performance on several datasets, they suffer from low robustness. For example, ASRS achieves 100% accuracy on dataset-23 (*Nutt-2003-v3*), but it yields even worse results than that of random assignment on dataset-9 (*Chen-2002*). We emphasize that, for unsupervised tasks, robustness is much more important than performance in practice when dealing with highly heterogeneous molecular data types (such as mRNA expression)(*16*). Different from LCE and ASRS, ECC fuses the basic partitions in a utility way, which ensures highly meaningful interpretations with high stability for the final consensus partition (**Supplementary Materials Sec. I**). To compare the overall performance of those clustering methods over the 35 benchmark datasets, we proposed an average performance score (see **Materials and Methods**) and found that ECC revealed significant advantages over all other methods in terms of average performance score. We notice that there are four specific datasets (*Gordon-2002, Khan-2001, Ramaswamy-2001* and *Shipp-2002*) for which all clustering methods yield very poor performance, most likely due to the presence of irrelevant or noisy features. We pointed that this difficulty cannot be easily resolved by any existing clustering methods. Yet, it can be alleviated by a complementary basic partition generation strategy of RPS, i.e., the *Random Feature Selection* (RFS) strategy, within the framework of ECC. To achieve that, we generate different sub-datasets by randomly selecting certain percentage of features (e.g., mRNAs) and then apply traditional clustering (e.g., *K*-means) to those sub-datasets to obtain basic partitions. Indeed, we find that for these four datasets, the performance of RFS exceeds RPS with all sampling ratios. This indicates that RFS helps us avoid noisy and irrelevant mRNA expressions (see **Supplementary Materials Sec. IV** for details).

**Fig. 2.**
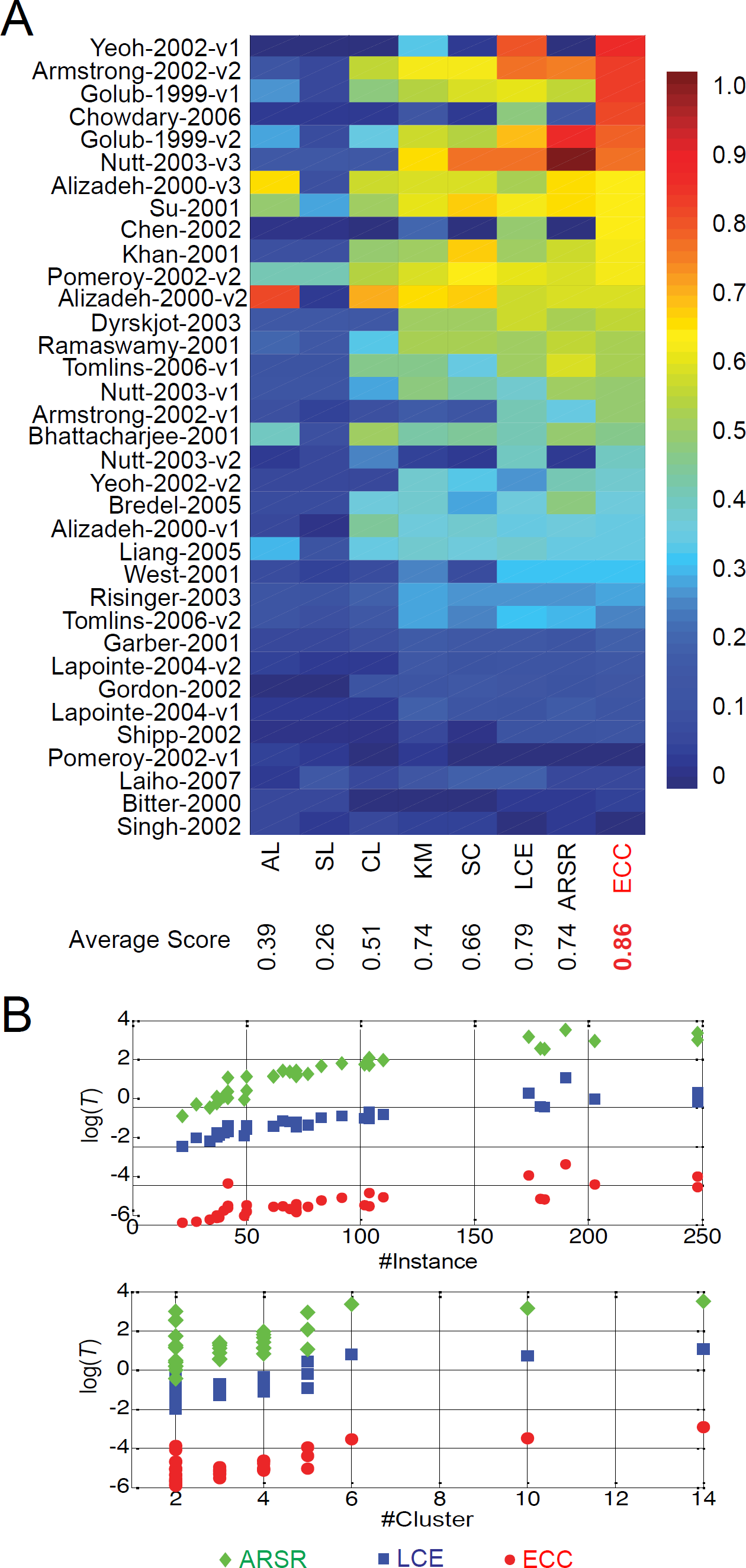
The performance of ECC on 35 benchmark cancer gene expression datasets. (**A**), The performance of different clustering methods (five traditional clustering methods: Agglomerative Hierarchical Clustering with Average Linkage (AL), Single-Linkage (SL) and Complete-Linkage (CL), *K*-means Clustering (KM), and Spectral Clustering (SC); and two state-of-the-art consensus clustering methods: the Link-based Cluster Ensemble (LCE) and Approximate SimRank-based (ASRS) methods) is measured by the Normalized Mutual Information (*NMI*). Overall, ECC outperforms the traditional clustering methods and state-of-the-art consensus clustering methods by a large margin. (**B**), The execution time *T* (in logarithmic scale) of different consensus clustering methods (ARSR, LCE, and ECC) as a function of the number of instances or the number of classes.

In addition, ECC has tremendous merits in terms of computational cost. **Fig. 2B** shows the execution time (in logarithmic scale) of the three consensus clustering methods (LCE, ASRS and ECC). The time complexity of ECC is *O*(*InKr*), where *I* is the number of iterations, *n* is the number of subjects, *K* is the number of clusters and *r* is the number of basic partitions. The space complexity of ECC is *O*(*nr*). For LCE and ASRS, the space complexities are both *O*(*n*^2^); and the time complexities are *O*(*n*^2^log*n*) and *O*(*n*^3^), respectively. Naturally, ECC is more suitable for high-throughput molecular data analysis (**Supplementary Materials Sec. IV**). For example, on dataset-34 (*Yeoh-2002-v1*), ECC is 115 times and 1,600 times faster than LCE and ASRS, respectively.

### Translational applications of ECC

The availability of massive and various molecular data types generated from large-scale and well-characterized cohorts across multiple cancer types provides an unprecedented opportunity for patient stratification. Here we demonstrated the translational applications of ECC based on13 major cancer types from The Cancer Genome Atlas (TCGA) project with sufficient sample size and clinical profiles for four molecular data types: mRNA expression (RNA-seq V2), microRNA (miRNA) expression, protein expression, and somatic copy number alterations (SCNAs), as shown in **Supplementary Table S3**. For fair comparison, we collected the empirical number of clusters (subtypes) for the 13 TCGA cancer types from previous studies. Then we applied survival analysis to evaluate the performance of different clustering methods in terms of −log_10_(*P*) with *P* the log-rank test *P*-value (**Supplementary Materials Sec. V** and **Supplementary Tables S7-10**).

For each molecular data type (Protein, miRNA, mRNA, and SCNA), we calculated the clustering performance of ECC against other clustering methods across the 13 TCGA cancer types. We found that ECC outperformed other methods (in terms of the number of significant survival analysis results across the 13 TCGA cancer types, as highlighted in dotted red rectangles in **Fig. 3A-D**) for any single molecular data type.

**Fig. 3.**
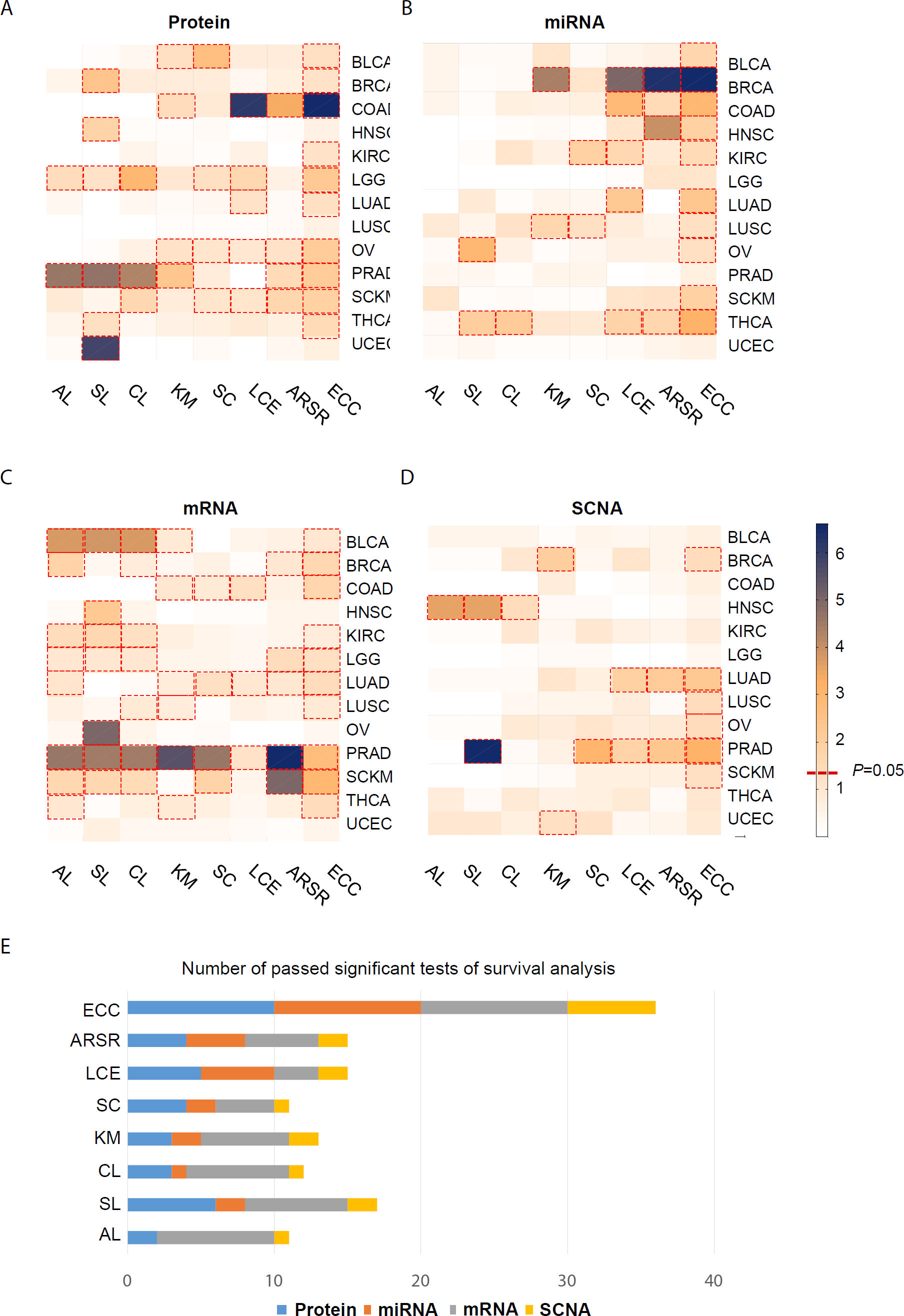
Performance of 7 different clustering methods on 4 molecular data types across 13 major cancer types from TCGA. Heatmaps show the survival analysis for 13 major cancer types using 7 different clustering methods based on four molecular data types: (**A**) protein expression (protein), (**B**) miRNA expression (miRNA), (**C**) mRNA expression (mRNA), and (**D**) somatic copy number alterations (SCNA), respectively. We use the −log(*P*) to draw the heatmap and elements with dotted red rectangles have *P* <0.05. (**E**), This plot displays for each clustering method the times that it passes the significant tests of survival analysis, i.e. the number of dotted red rectangles in (**A-D**), over the 13 cancer types and the 4 different molecular data types.

By integrating the 4 different molecular data types, ECC generated significant clusters (cancer subtypes) for all the 13 TCGA cancer types (*P* < 0.05, log-rank test, **Table. 1**). Note that traditional clustering methods and existing consensus clustering methods cannot easily integrate multiple molecular data types, due to the presence of missing values for certain molecular data type of certain subjects. Yet, ECC can naturally resolve this issue by utility fusion, where missing values in basic partition provide no utility for the final fusion (**Supplementary Fig. S9**). Moreover, by integrating multiple molecular data types, ECC is effectively more robust to noise present in the data (partially because it has more data types to generate basic partitions). For example, in the case of uterine corpus endometrial carcinoma (UCEC), using any of the 4 molecular data types, ECC cannot yield significant clusters (**Fig. 4A**). Yet, by integrating multiple molecular data types (pan-omics), ECC yielded 4 significant clusters (**Fig. 4B**) with distinct patient survival curves (*P* = 0.0043, **Fig. 4C**); while using any single molecular data type the clusters generated by ECC do not pass the significance test of survival analysis (*P* >0.05, **Supplementary Tables S7-10**). In addition, subtypes identified by ECC via integrating 4 molecular data types were closely associated with the clinical subtypes on a histological basis in UCEC (**Fig. 4D**). For instance, subtype 2 with most aggressive uterine tumor shows poor survival than subtype 1 with the less aggressive uterine tumors. Similar trends are also observed in ovarian serous cystadenocarcinoma (OV, *P* = 7.79×10^−4^) and prostate adenocarcinoma (PRAD, *P* = 5.27×10^−4^) (see **Supplementary Table S11**). Since TCGA clinical information may not be complete or rigorously annotated, future efforts of assessing the clinical utility of different subtypes on additional patient cohorts with more carefully annotated clinical variables are needed.

**Table. 1.**
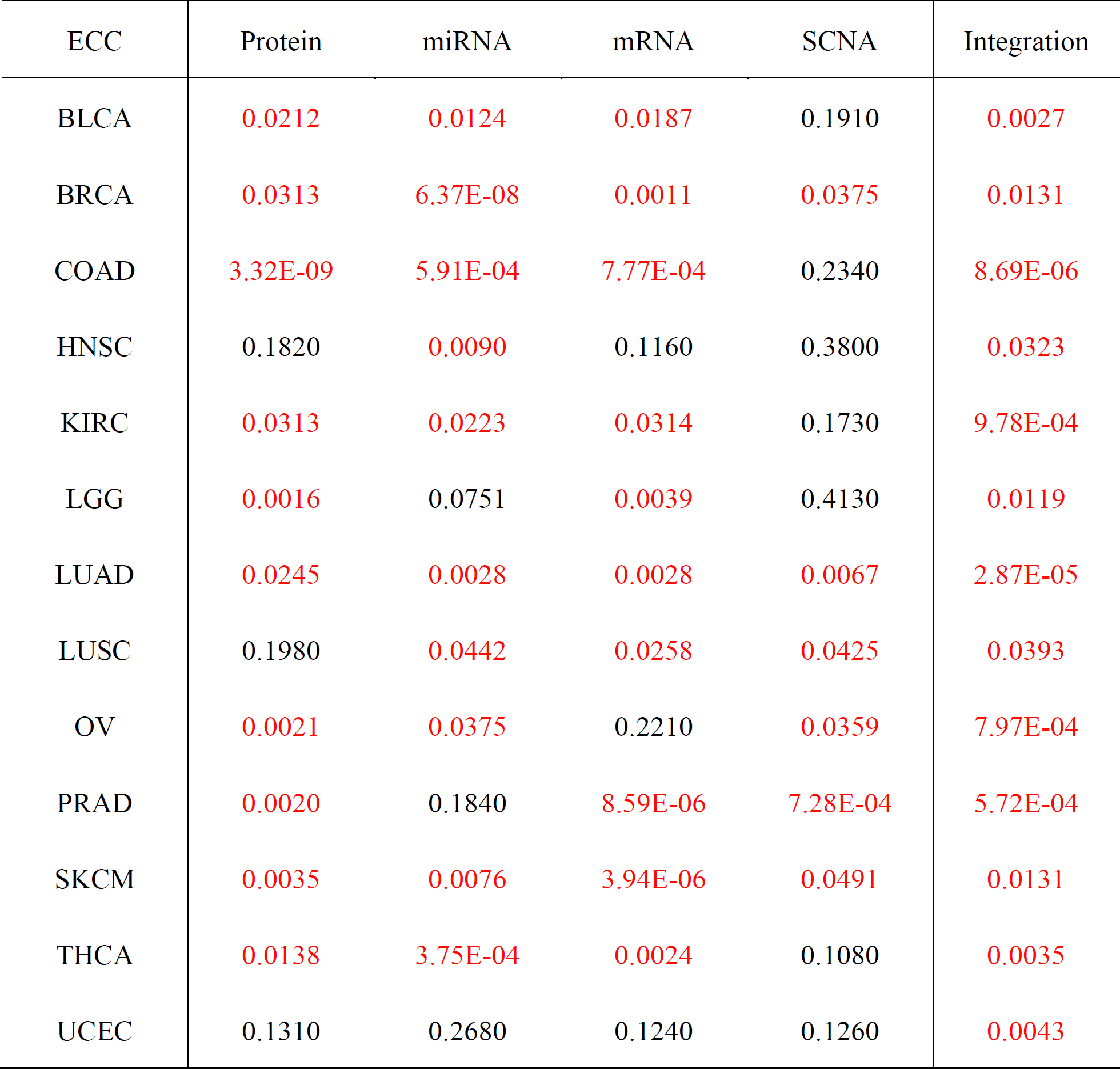
Performance of ECC on 4 molecular data types and its integration across 13 major cancer types from TCGA. The performance is quantified by the log-rank test *P*-value of the survival analysis over the identified clusters (cancer subtypes). We highlight *P* <0.05 in red. With the integration of the 4 molecular data types, i.e., the pan-omics, ECC yields clusters that pass the significant test for all the 13 cancer types.

| ECC | Protein | miRNA | mRNA | SCNA | Integration |
| --- | --- | --- | --- | --- | --- |
| BLCA | 0.0212 | 0.0124 | 0.0187 | 0.1910 | 0.0027 |
| BRCA | 0.0313 | 6.37E-08 | 0.0011 | 0.0375 | 0.0131 |
| COAD | 3.32E-09 | 5.91E-04 | 7.77E-04 | 0.2340 | 8.69E-06 |
| HNSC | 0.1820 | 0.0090 | 0.1160 | 0.3800 | 0.0323 |
| KIRC | 0.0313 | 0.0223 | 0.0314 | 0.1730 | 9.78E-04 |
| LGG | 0.0016 | 0.0751 | 0.0039 | 0.4130 | 0.0119 |
| LUAD | 0.0245 | 0.0028 | 0.0028 | 0.0067 | 2.87E-05 |
| LUSC | 0.1980 | 0.0442 | 0.0258 | 0.0425 | 0.0393 |
| OV | 0.0021 | 0.0375 | 0.2210 | 0.0359 | 7.97E-04 |
| PRAD | 0.0020 | 0.1840 | 8.59E-06 | 7.28E-04 | 5.72E-04 |
| SKCM | 0.0035 | 0.0076 | 3.94E-06 | 0.0491 | 0.0131 |
| THCA | 0.0138 | 3.75E-04 | 0.0024 | 0.1080 | 0.0035 |
| UCEC | 0.1310 | 0.2680 | 0.1240 | 0.1260 | 0.0043 |

**Fig. 4.**
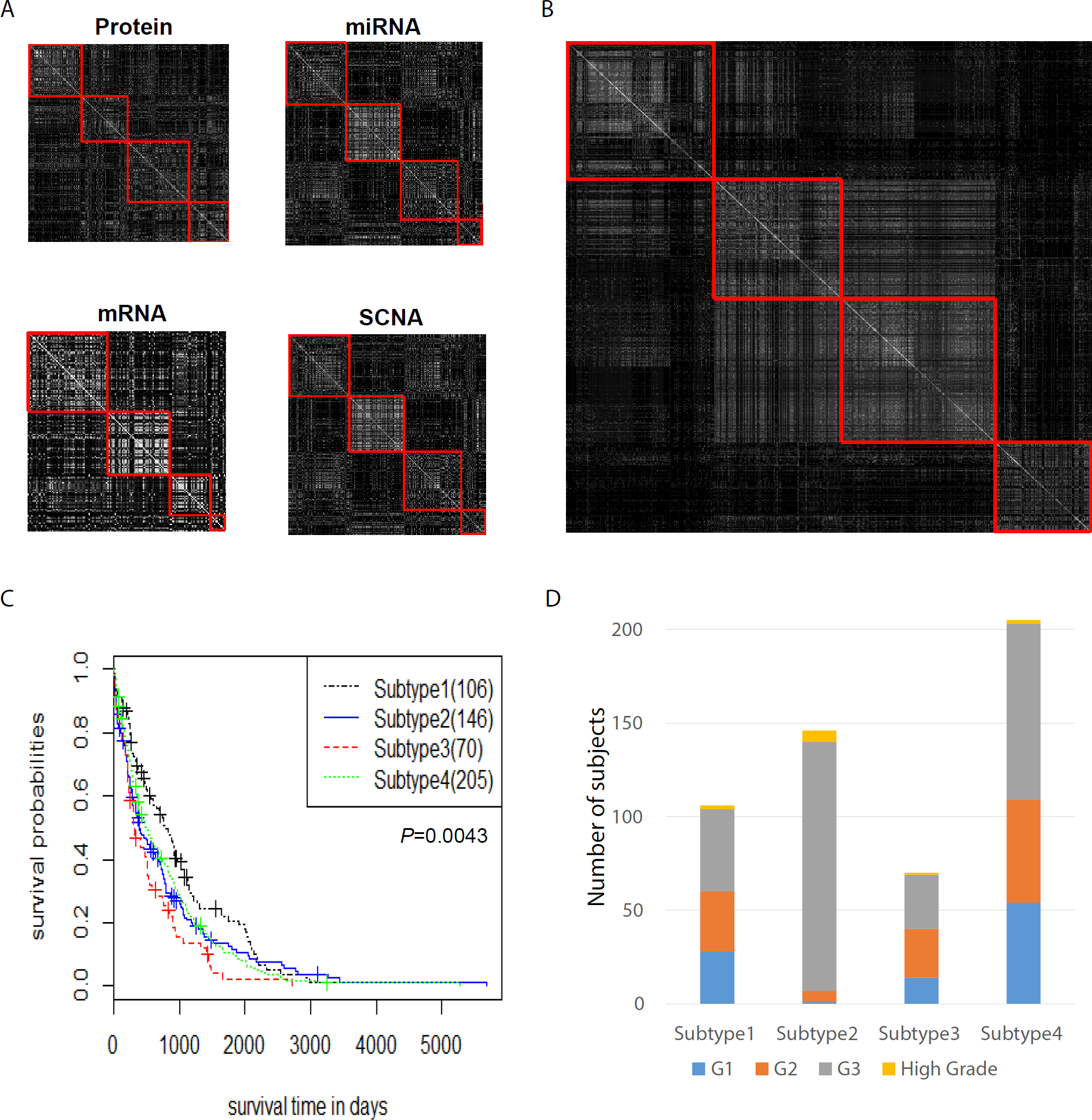
Performance of ECC foruterine corpus endometrial carcinoma (UCEC) subjects from TCGA. The similarity matrices calculated from the 4 clusters generated by ECC using single molecular data type (**A**) and pan-omics data (**B**) of UCEC. The survival curves (**C**) and the composition of different clinical subtypes (**D**) for the 4 clusters generated by ECC using pan-omics data of UCEC.

## Discussion

In sum, we show that ECC owns significant advantages in terms of cluster validity, execution time and space complexity, and robustness compared with other clustering methods in patient stratification. We demonstrate that ECC with RFS strategy can alleviate the detriment effect of irrelevant and noisy features. Moreover, ECC displays superior performance on the pan-omics data by integrating multiple molecular data types than that of single molecular data type. We anticipate that integrating more types of both molecular and clinical data, such as somatic mutations, DNA methylation, functional genomic data generated from CRISPR/Cas9 (*17*), proteogenomics (*18*), radiomics (*19*), and electronic medical records (*20*), will further improve patient stratification. Altogether, our ECC method paves the way to a much more refined representation and understanding of various molecular data types, facilitating the development of precision medicine.

## Methods

### Consensus clustering

Here we introduce the basic ideas of consensus clustering in the context of omics data (e.g., gene expression) analysis. Consensus clustering was originally developed for fusing several existing partitions into a robust one (*1*), and has recently been applied to gene expression data analysis (*2*,*3*). For example, the link-based cluster ensemble (LCE) method first summarizes several basic partitions into a co-association matrix (that measures how often two instances simultaneously occur in the same cluster); then modifies the zero entries in the co-association matrix with the distance derived from the original data; and finally conducts spectral clustering to obtain the consensus partition (*2*). As a variant of the LCE method, the Approximate SimRank-based(ASRS) method employs very similar idea with slightly different modification on the zero entries in the co-association matrix (*3*).

Different from the existing consensus clustering methods (LCE and ASRS), our ECC method employs an entropy-based utility function for the guidance of fusing all the basic partitions into a consensus one, which has a more meaningful interpretation than existing consensus clustering methods. Let X denote a gene expression dataset with *n* subjects and *m* genes. A partition of X into *K* crisp clusters is represented as a collection of *K* subsets of instances in C = {C_k_ | *k* = 1,…,*K*}, with 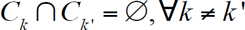, and 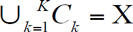 or as a label vector *π =*(*L_π_*(*x_1_*),*…*,*L_π_*(*x_n_*))^T^ where *L_π_* maps *x_l_* to some label in {1,2,…,*K*}, 1 ≤ *l* ≤ n. Suppose we have *r* basic partitions denoted as{*π*^(1)^,*π*^(2)^,…,*π*^(*r*)^} generated by some traditional clustering method (e.g., *K*-means) and there are *K_v_* clusters in *π*^(*v*)^, for 1 ≤ *v* ≤ *r*. The goal of consensus clustering is to find a consensus partition *π* by solving the following optimization problem:

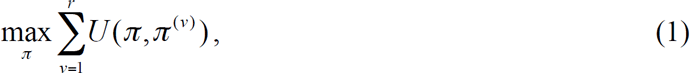

where *U* is a utility function measuring the similarity at the partition-level between each basic partition and the consensus one. In other words, we expect to find an optimal partition that agrees with the basic ones as much as possible. Different utility functions measure the similarity of two partitions in different aspects, rendering different objective functions for consensus clustering. In this work, an entropy-based utility function is employed for its fast convergence and high quality (*4*).

### Entropy-based Utility Function

The core of ECC is to fuse these basic partitions into a consensus one based on an entropy-based utility function, which assures the consensus clustering algorithm to be highly efficient and robust. As formulated in Eq. (1), a utility function is defined on two partitions *π* and *π*^(*v*)^ to measures their similarity at partition-level. We can employ the following contingency table to calculate the entropy-based utility function.

|  |  |  |  |  |  |  |
| --- | --- | --- | --- | --- | --- | --- |
| | | $\pi^{(v)}$ | | | | |
| | | $C'_1$ | $C'_2$ | ⋯ | $C'_{K_v}$ | $\Sigma$ |
| $\pi$ | $C_1$ | $n_{11}$ | $n_{12}$ | ⋯ | $n_{1K_v}$ | $n_{1+}$ |
| | $C_2$ | $n_{21}$ | $n_{22}$ | ⋯ | $n_{2K_v}$ | $n_{2+}$ |
| | · | · | · | ⋯ | · | · |
| | $C_K$ | $n_{K1}$ | $n_{K2}$ | ⋯ | $n_{KK_v}$ | $n_{K+}$ |
| | $\Sigma$ | $n_{+1}$ | $n_{+2}$ | ⋯ | $n_{+K_v}$ | $n$ |

Here we have two partition *π* and *π*^(*v*)^, which contain *K* and *K_v_* clusters, respectively. We assume *π* is the ground truth, and *π*^(*v*)^ is the clustering result generated by a specific clustering algorithm. Let *n_ij_* denote the number of objects shared by cluster *C_i_* in *π* and cluster *C_j_* in *π*^(*v*)^. Define 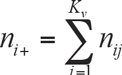, and 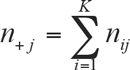, 1 ≤ *i* ≤ *K*, 1 ≤ *j* ≤ *K_v_*.

Based on the contingency table, for *π* and *π*^(*v*)^ we define two discrete distributions 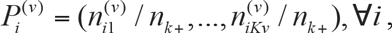, and 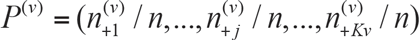. Then we have

#### Definition 1

(*U_H_*). An *entropy-based utility function U_H_* is defined as

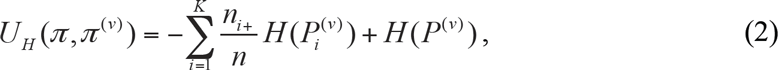

*where H denotes the Shannon entropy*.

Since Shannon entropy is a concave function, according to the Jensen’s inequality, we can prove that 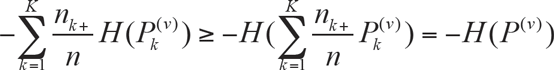, rendering that *U_H_* ≤ 0. A larger *U_H_* indicates the higher utility from the two partitions in greater similarity. Note that *U_H_* is asymmetric, with *U_H_*(*π*,*π*^(*v*)^) ≠ *U_H_*(*π*^(*v*)^,*π*), if *π*≠*π*^(*v*)^.

### Entropy-based Consensus Clustering

Although it is crucial to design a utility function, how to optimize it in an efficient way is another challenge. Thanks to the general *K*-means based Consensus clustering (*4*), which has substantial advantage in terms of efficiency; we can transform the optimization problem in Eq.(1) into a modified *K*-means clustering problem as follows.

Let **B** = (*b_1_,…,b_n_*)^T^ be a binary matrix derived from *r* basic partitions{*π*^(1)^,*π*^(2)^,…,*π*^(*r*)^}, with

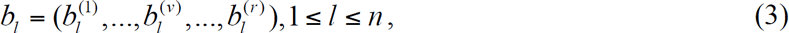

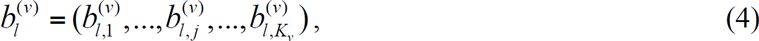

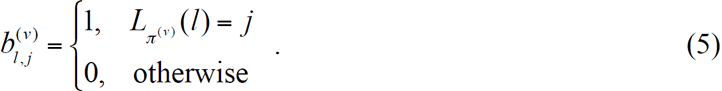

Apparently, **B** is a 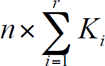 binary matrix, with 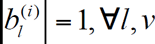 For the Entropy-based Consensus Clustering, a *K*-means clustering is directly conducted on **B** with the following modified distance function.

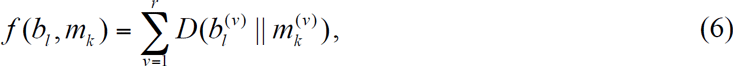

where 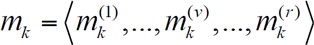 with 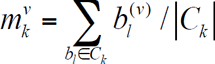, and 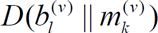 KL-divergence from *b_l_*^(*v*)^ to *m_k_*^(*v*)^.

By this means, the complex consensus clustering can be exactly mapped into a classic *K*-means clustering with a modified distance function, which has roughly linear time complexity and its convergence can also be guaranteed as well. The exactness of the mapping can be rigorously proved (see **Supplementary Materials Sec. I.A** for details). This mapping makes ECC very practical for large-scale molecular data analysis. Indeed only *r* elements are non-zero entries in each row of **B**, which leads the time complexity from 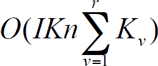 to *O*(*IKnr*), where *I* is the number of iterations.

### Handling missing values

Missing values are quite common in practice due to data collection or device failure, especially for the pan-omics data of a large population (in computer science, this kind of data is called multi-view). Typically there are two ways to handle those missing values. One is to just remove the instances (i.e., subjects) that have missing values in any single molecular data type (or any single view). Apparently, this is of great waste because those instances (subjects) might have values for many other views (molecular data types). The other way is to replace these missing values by default or average values. This would harm the original data structure and degrade the clustering performance. We can naturally resolve this challenging issue within the framework of ECC. In particular, we consider that those missing values, which lead to missing labels in the basic partitions, do not provide any utility for the consensus fusion. If a basic partition has missing labels, we call it an *incomplete basic partition* (IBP). For IBP, we directly denote 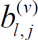 as an all-zero vector, which will not be involved in the distance calculation and centroid update. The following is the distance function for IBP:

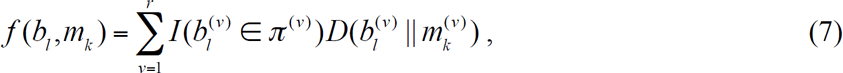

and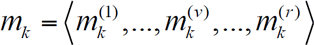 with

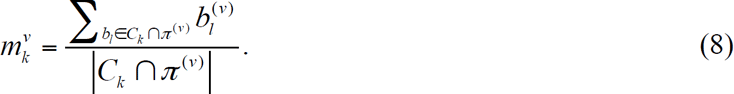

### Datasets

In this work, we use 110 synthetic datasets to systematically evaluate the performance of ECC. The 110 synthetic datasets are generated by a well-established dynamical gene regulation model (*5*):

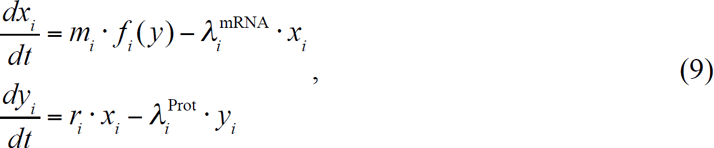

where *m_i_* is the maximum transcription rate, *r_i_* is the translation rate, 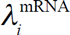 and 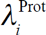 are the mRNA and protein degradation rates, and **x** and **y** are vectors of mRNA and protein concentration levels, respectively. *f_i_*(·) computes the relative activation of gene. The topology of the gene regulatory network is encoded in the activation functions.

Among the 110 synthetic datasets, 55 of them are based on an Erdős–Rényi random network with 500 nodes (genes), and the other 55 are based on a human transcriptional regulation network of 2723 genes (*6*). Each dataset contains 200 subjects with its benchmark of 4 clusters (50 subjects in a cluster). Each dataset contains 200 subjects divided evenly into 4 groups (clusters). Each group has a specific set of knocked-out genes. A more detailed description of the synthetic datasets can be found in **Supplementary Materials Sec. I**.

Besides the 110 synthetic datasets, 35 widely used cancer gene expression benchmark datasets (*7*) are employed to test the cluster validity of ECC. Also, 13 cancer types with four molecular data types from TCGA with survival information are used for practical evaluation of ECC (**Supplementary Table S3**).

### Evaluation metrics

Since the true labels for synthetic and benchmark datasets are available, we can apply external measurements to objectively evaluate the performance of different clustering algorithms. Although there are many external measurements, some of them are biased. According to Wu *et.al.* (*12*), two normalized external measurements, *NMI* and *R_n_* are unbiased and hence can be chosen for proper evaluation of clustering performance. Both can easily be calculated from the contingency table.

Normalized Mutual Information (*NMI*) measures the mutual information between resulted cluster labels and ground truth labels, followed by a normalization operation to assure *NMI* ranges from 0 to 1. Mathematically, it is defined as:

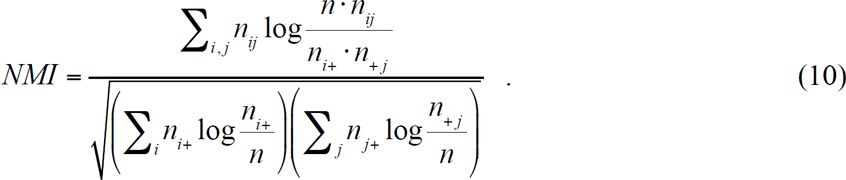

Normalized Rand Index, denoted as *R_n_* measures the similarity between two partitions in a statistical way, which is defined as:

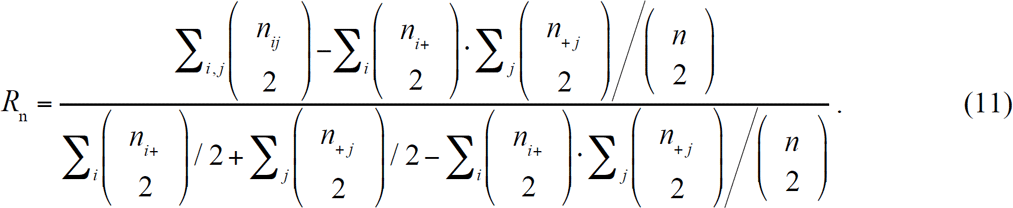

Note that both *NMI* and *R_n_* are positive measurements, i.e. a better partition has a larger *NMI* or *R_n_* value. Although *R_n_* is normalized, it can still be negative, which means that the partition is even worse than random label assignment.

To compare the overall performance of those clustering algorithms over the 35 benchmark cancer expression datasets, we propose an average performance score as follows:

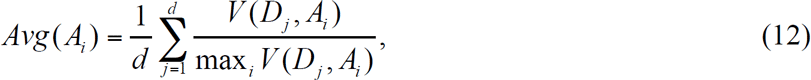

where*V*(*D_j_,A_i_*) denotes the performance (i.e., *R_n_* or *NMI*) of Algorithm *A_i_* on dataset *D_j_* and *d* is the total number of benchmark datasets.

### Code availability

The MATLAB code is freely available at http://scholar.harvard.edu/yyl/ecc.

## Acknowledgments

We thank Rulla Tamimi and Edwin Silverman for valuable discussions. This work is supported in part by the John Templeton Foundation (Award number 51977), the National Academy of Sciences-Grainger Foundation Frontiers of Engineering Award (2000006959), NSF CNS Award (1314484), ONR award (N00014-12-1-1028), ONR Young Investigator Award (N00014-14-1-0484), and U.S. Army Research Office Young Investigator Award (W911NF-14-1-0218).

## Contributions

Y.-Y.L. and Y.F. conceived the project. Y.-Y.L. designed the research. H.L. developed the code to perform ECC clustering and conducted all the cluster analyses. H.F. and R.Z developed the code to generate synthetic gene expression data. R.Z. collected cancer gene expression benchmark datasets. F.C. collected TCGA datasets and interpreted results. All authors analyzed the results. F.C., H.L., R.Z. and Y.-Y.L. wrote the manuscript. H.F. edited the manuscript.

## Author Information

The authors declare no competing financial interests. Correspondence and requests for materials should be addressed to F.Y. or Y.-Y.L.

## Supplementary Information

This file is the supplementary information for our paper “A Novel Clustering Algorithm for Patient Stratification”, which contains the theoretical analysis of our ECC method, the generation of synthetic datasets, the description of real datasets, and additional numerical results.

### I. THEORETICAL ANALYSIS OF ECC

In this work, we map the complicated utility optimization problem in ECC to a classic *K*- means clustering problem with a modified distance. Here we rigorously prove the correctness and convergence of ECC. We summarize all the variables used in this section in Table SI.

### A. Correctness of ECC

To prove the correctness of ECC, we consider the following contingency matrix.

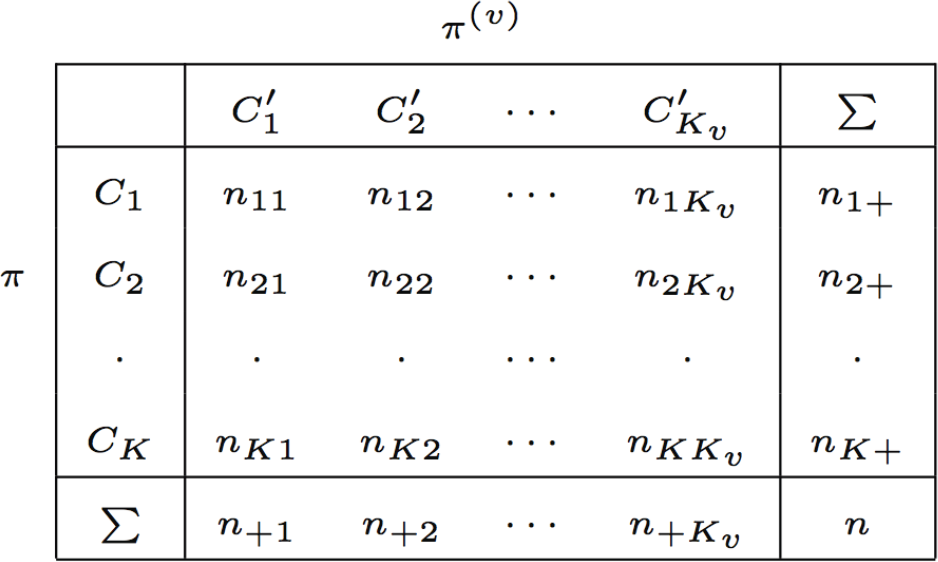

| | | $\pi^{(v)}$ | | | | |
| --- | --- | --- | --- | --- | --- | --- |
| | | $C'_1$ | $C'_2$ | ⋯ | $C'_{K_v}$ | $\sum$ |
| $\pi$ | $C_1$ | $n_{11}$ | $n_{12}$ | ⋯ | $n_{1K_v}$ | $n_{1+}$ |
| | $C_2$ | $n_{21}$ | $n_{22}$ | ⋯ | $n_{2K_v}$ | $n_{2+}$ |
| | · | · | · | ⋯ | · | · |
| | $C_K$ | $n_{K1}$ | $n_{K2}$ | ⋯ | $n_{KK_v}$ | $n_{K+}$ |
| | $\sum$ | $n_{+1}$ | $n_{+2}$ | ⋯ | $n_{+K_v}$ | $n$ |

Here we have two partition *π* and *π*^(*v*)^. We assume *π* is the ground truth, and *π*^(*v*)^ is the clustering result returned by a certain clustering algorithm. Let *n_ij_* denote the number of data objects shared by both cluster 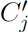 in *π*^(*v*)^ and cluster *C_i_* in *π*, 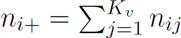 and 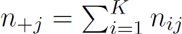 1 ≤ *i* ≤ *K*,1 ≤ *j* ≤ *K_v_*,

Let **B** = (*b*_1_,…,*b*_n_)^⊤^ be a binary matrix derived from *r* basic partitions *π*^(1)^,…,*π*^(r)^, with

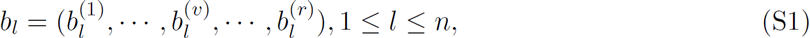

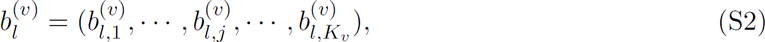

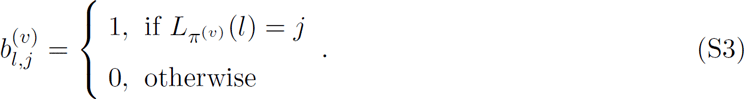

Apparently, **B** is a 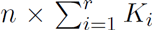 binary matrix, with 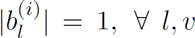 For the Entropy-based Consensus Clustering, a K-means clustering is directly conducted on **B** with the modified distance function defined as follows.

#### Theorem 1.

*Assume *π* is a consensus partitioning of X, with K clusters C_1_,…,C_K_. Given r basic partitions *π*_1_,…,*π*_r_, we have*

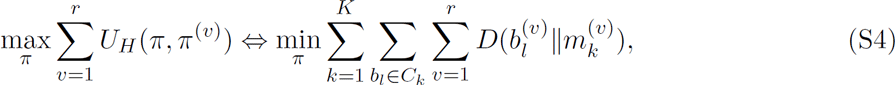

*where 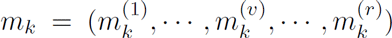 with 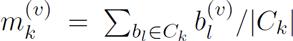, 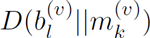 is the KL-divergence from 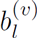 to 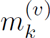 and U_H_ is the entropy-based utility function, which can be calculated as follow,*

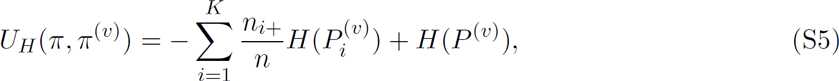

*where 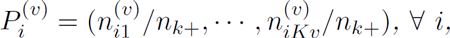, and 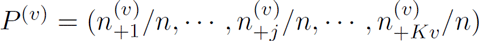.*

*Proof.* According to Bregmen Divergence[3], we have that 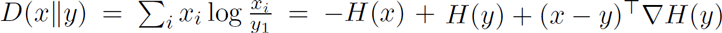, where 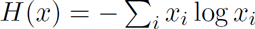 is the Shannon entropy. Hence we have

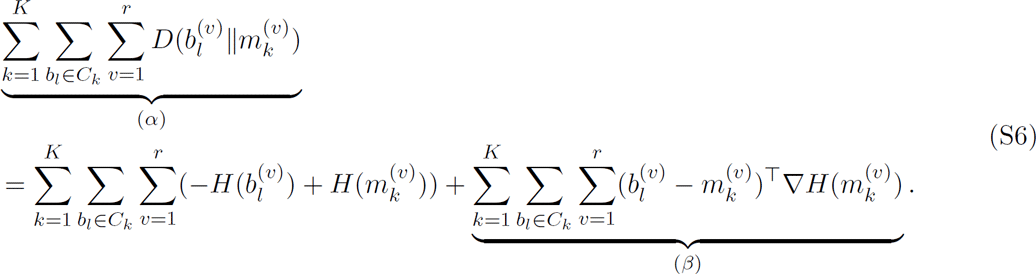

Since 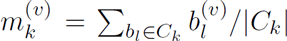, we have 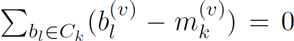,which indicates that the term (*β*) = 0. We therefore have:

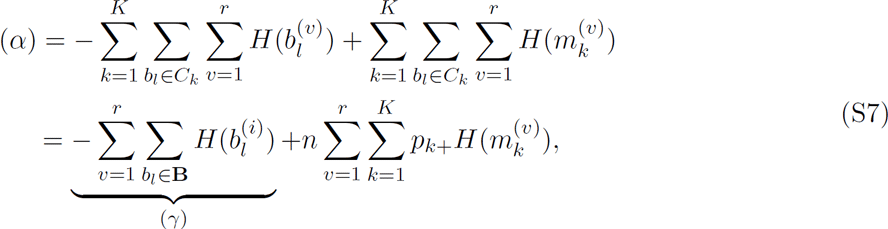

where *p_k+_/n*. Since the term (*γ*) and *n* are constants, we have

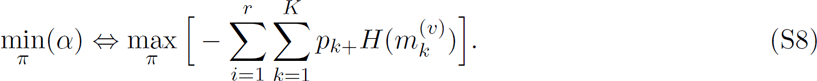

Note that 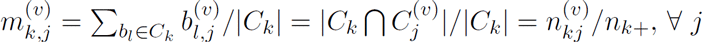, which indicates that

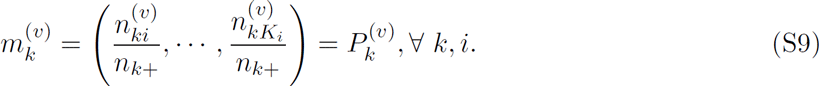

If we substitute *m_k_*^(*v*)^ by *P_k_^v^* in Eq. S*8*, and add the constant 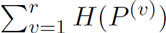 to the right-hand-side, we finally have

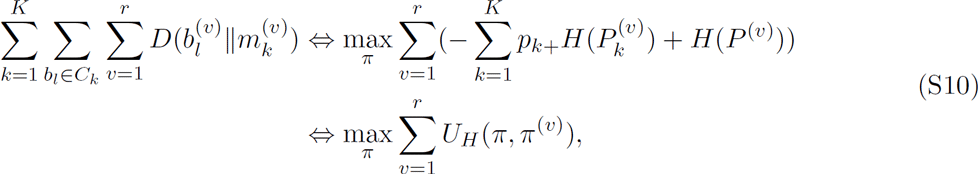

and the theorem thus follows.

#### Remark 1.

*Theorem 1 gives a new insight of the objective function. ECC aims to find a partition that agrees with the basic ones as much as possible and employs U_H_ to measure the similarity of two partitions. Theorem 1 ensures that we can calculate the distance of two partitions by KL-divergence to achieve the same goal.*

#### Remark 2.

*By Theorem 1, we can solve the consensus clustering problem by the classic K-means clustering, which is the fastest clustering algorithm. Recall that only r elements are non-zero entries in each row of **B**, thus the positions for these non-zero elements are needed, rendering the time complexity O(IKnr), where I is the iteration number. Usually, I,K,r are smaller than n. Therefore, the time complexity of ECC is roughly linear to the number of instances, which is suitable for high-throughput molecular data analysis.*

### B. ECC Algorithm

The pseudo code of our ECC algorithm is shown in Algorithm 1. In essence, ECC is a variant of K-means, which has the two-phase iteration: instance assignment and centroid update. The only difference between ECC and K-means is the distance function. In K-means, the squared Euclidian distance is employed, while we use a summation of several KL-divergence in ECC.

ECC has tremendous merits in terms of efficiency over the other methods. Fig. S8 shows the execution time (in logarithmic scale) of three consensus clustering methods (LCE, ASRS and ECC). The time complexity of ECC is *O*(*InKr*), where *I* is the iteration number, *n* is the subject number, *K* is the class number and *r* is the number of basic partitions. The space complexity of ECC is *O*(*nr*). For LCE and ASRS, the space complexities are both *O*(*n*^2^) and the time complexities are *O*(*n*^2^log*n*) and *O*(*n*^3^), respectively.

#### Algorithm 1

The algorithm of Entropy-based Consensus Clustering

~~~
**Input:** *X: data matrix, n×m;*
  *K*: number of clusters;
  *r*: number of basic partitions.
**Output:** Partition *π*;
1: Obtain the set of basic partitions n by some generation strategy;
2: Build the binary matrix *B* by Eq.(6-8) in the main paper;
3: Randomly select *K* instances from *B* as centroids;
4: **repeat**
5: Assign each instance to its closest centroid by the distance function in Eq.(9) in the main paper;
6: Update centroids by arithmetic mean;
7: **until** *K* centroids remain unchanged.
8: Return the partition *π*.
~~~

### C. Convergence of ECC

ECC is solved by the *K*-means clustering with a modified distance function. The convergence of ECC is assured by the following theorem [4].

#### Theorem 2.

*For the objective function in Theorem 1, ECC is guaranteed to converge in finite two-phase iterations of K-means clustering.*

*Proof.* The *K*-means distance function can be generalized as the Bregman divergences[3],

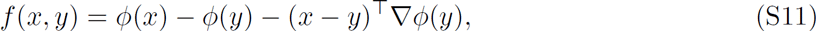

where *ϕ*(·) is a convex function. By using Bregman divergences, the convergence of K-means is guaranteed [3].

For the objective function in Theorem l, we have

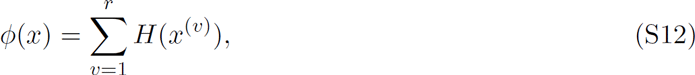

which is the summation of the Shannon Entropy. Since *H*(·) is a convex function and the summation preserves the convex property, therefore the distance function in Theorem 1 is a Bregman divergence, the convergence of ECC can then be guaranteed.□

In handling IBPs, the convergence property of ECC still holds.

## II. SYNTHETIC DATASETS

The 110 synthetic gene expression datasets are generated through a well-known dynamical gene regulation model[1]. To be self-contained, we summarize all the variables used in this section in Table SII..

### A. Dynamical Gene Regulation Model

This model is based on a gene regulatory network represented by a digraph *G*(*V,E*). Here*V* is the set of nodes (genes) and *E* is the set of directed edges (gene regulations). Both transcription and translation are modeled using a standard thermodynamic approach[1]. For each node *v_i_, i=1,2,…,n*, the changing rate of mRNA concentration *F_i_^mRNA^* and the changing rate of protein concentration *F_i_^Prot^* are described by a set of coupled ordinary differential equations (ODEs):

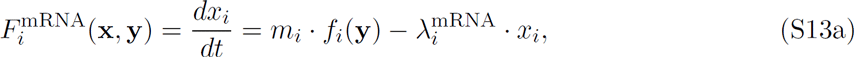

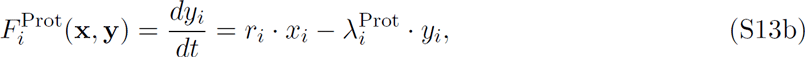

where *m_i_* is the maximum transcription rate, *r_i_* is the translation rate, λ_*i*_^mRNA^ and λ_*i*_^Prot^ are the mRNA and protein degradation rates, **x** ∈ **R**^n^ and **y** ∈ **R**^n^ are vectors of mRNA and protein concentration levels, respectively. *f_i_*(*·*) is the activation function of gene *i*, which is between 0 (gene *i* is turned off) and 1(gene *i* is maximally activated) given the protein concentrations **y**. The network topology is encoded in the activation functions. A more detailed definition of *f_i_*(·) is as follows.

We use a standard thermodynamics-based approach to model gene regulation[1]. The basic assumption is that binding of transcription factors (TFs) to cis-regulatory sites on the DNA is in quasiequilibrium because it is orders of magnitude faster than transcription and translation. In the simplest cases, gene *i* is regulated by a single TF (e.g.*TF_j_*), then its promoter has only two states: either the *TF_j_* is bound (state *S*_1_) or un-bound (state *S*_0_). The probability *P*(*S*_1_) that gene *i* is in state *S*_1_ at a certain time instant is given by the fractional saturation:

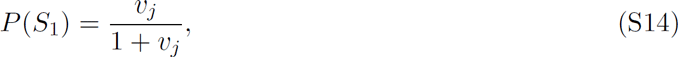

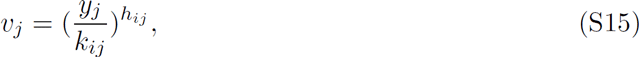

where *y_j_* is the concentration of *TF_j_*, *k_ij_* is the dissociation constant, and *h_ij_* the Hill coefficient. The bound TF activates or represses the expression of the gene. In state *S_0_*, the relative activation is*α_0_*; in state S_1_, the relative activation is *α_1_*. Given *P*(*S*_1_) and its complement *P*(*S*_0_) = 1 — *P*(*S*_1_), we can derive the function *f_i_(y_j_)*), which computes the mean activation of the gene *i* as a function of the TF concentration *y_j_*:

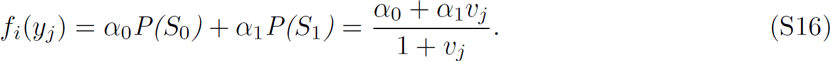

We can also consider gene *i* has two regulatory inputs. The resulting expression would be:

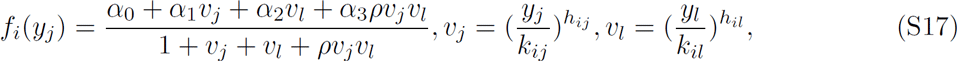

where *ρ* is the cooperativity factor, and *α*_i_ are the relative activations when none of the TFs (*α*_0_), only the first (*α*_1_, only the second (*α*_2_) or both TFs are bound (*α*_3_).

This approach can be used for an arbitrary number of regulatory inputs. If a gene is regulated by *N* TFs, it will have *2^N^* states: each of the TFs can be bound or un-bound. Thus, the function for *N* regulators would be:

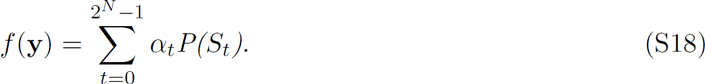

Based on thermodynamics, we can compute the probability *P*(*S_t_*) for every state *t*.

Assume now we have the gene regulatory network *G*(*V, E*) and the detailed dynamical model parameters. The initial values of *x_i_* and *y_i_* are generated randomly from the interval [0.001, 1]. Then we can compute the time evolution of **x** and **y** until the system reaches a steady state.

### B. Noise

The integration of the coupled ODEs (Eq.S1a and Eq.S1b) results in noiseless mRNA and protein concentration levels. However, both molecular and measurement noise in gene expressions are unavoidable in practice. In living cells, molecular noise originates from thermal fluctuations and stochastic processes such as transcription and translation. Moreover, measurement noise of gene expression depends on the experimental technology used to monitor the gene expression level.

### 1. Molecular Noise

Both *F_i_*^mRna^ and *F_i_*^Prot^ can be written as follows:

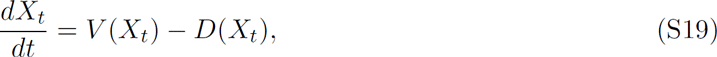

where *V*(*X_t_*) is the production term and *D*(*X_t_*) is the degradation term of mRNA or protein. To model molecular noise in the transcription and translation processes, we can use the following chemical Langevin equation (CLE):

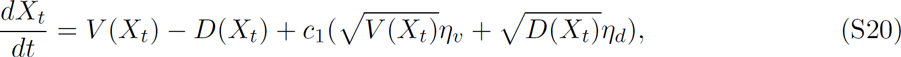

where *η_v_* and *η_d_* are independent Gaussian white-noise processes, *c_1_* is a constant to integrate two equations to control the amplitude of the molecular noise.

To solve the CLE, we can use the Stratonovich scheme and the Milstein method. Stratonovich Scheme is a technique used in stochastic integral, which is very similar to *Ito* integral. Suppose we have a stochastic differentiable equation:

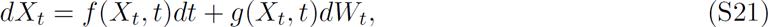

where *X_t_* is the random variable, *f* is the drift coefficient, *g* is the diffusion coefficient, and *W_t_* is the Wiener Process. Integrating both sides yields:

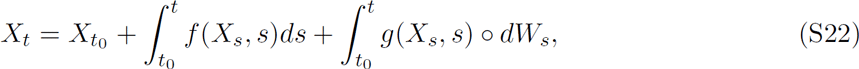

where ○ is used here to distinguish Stratonovich from*Ito*.

We compute the integrals in Eq. S10 using the Milstein method. Milstein method has two versions, namely a normal one and a derivative-free version. For simplicity, we use derivative-free Milstein method:

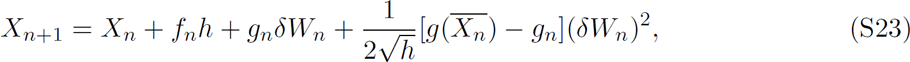

where

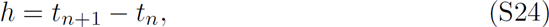

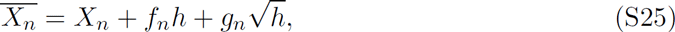

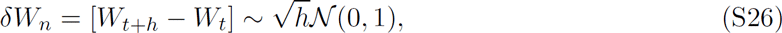

Then we update *X* until it reaches a steady state.

### 2. Measurement Noise

The measurement noise depends on the technology used to monitor level of gene expression and hence is modeled independently of the molecular noise. In this work, we model the measurement noise as follows:

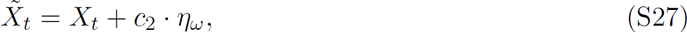

where *X_t_* is computed from Eq.S1, *η_ω_* represents independent Gaussian white-noise processes, and *c_2_* is a constant quantifying the measurement noise level.

### C. Synthetic Disease Subtypes

In order to generate synthetic disease subtypes, we first generate a random digraph *G* to represent the gene regulatory network. Considering the exponential time complexity of calculating the activation function, we set the average in-degree *k_in_* = 2 in the digraph *G*. Regarding the model parameters in Eq.S1, we choose *m_i_* and *r_i_* randomly from the interval [0, 1]. The Hill coefficient *h_ij_* is sampled from a Gaussian distribution *N*(2,4) bounded in the interval [1, 10]. An example of the gene regulatory network of 20 genes is shown in Fig. S1**a**. The time evolution of mRNA concentration levels *x_i_*(*t*) without noise, with only molecular noise, and with both molecular and measurement noise, are shown in Fig. S1**b**, Fig. S1**c**, Fig. S1**d**, respectively. This serves as the baseline model.

To simulate gene expression data for different disease subtypes, we assume that different disease subtypes are associated with different sets of genes that are knocked out. For those knock-out genes, we set their transcription rates *m_i_* to be zero. Suppose in total we have *N* subjects divided into four groups (*G*_0_, *G*_1_, *G*_2_, *G*_3_) evenly: subjects in *G*_0_ are generated from the baseline model, while subjects in *G*_1_, *G*_2_ and *G*_3_ have different sets of knock-out genes (see Fig. S2). For each group (disease subtype), different subjects are simulated from different initial conditions of **x** and **y**.

## III. REAL-WORLD DATASETS

We analysed 35 benchmark cancer gene expression datasets[2] with label (i.e. cluster structure) information to fully evaluate the performance of ECC, as well as 13 real cancer gene expression datasets with survival information available for practical evaluation. Some key characteristics of these datasets are summarized in Table SIII and Table SIV.

### A. Benchmark Gene Expression Datasets

For the 35 benchmark datasets in Table SIII, the numbers of subjects vary from 22 to 248, the numbers of genes vary from 85 to 4,553 and the numbers of clusters vary from 2 to 14. Some datasets are from the same source. For example, dataset 2 and 3 (*Alizadeh-2000-v2, Alizadeh-2000-v3*) share the same gene expression data, but with different cluster numbers; dataset-4 and 5 (*Armstrong-2002-v1, Armstrong-2002-v2*) have the same subject number, but different dimensions of gene expression; dataset-14 (*Golub-1999-v2*) splits one cluster in dataset-15 (*Golub-1999-v1*) into two; dataset-32 (*Tomlins-2006-v2*) has one more cluster than the dataset-33 (*Tomlins-2006-v2*).

### B. Molecular data from TCGA

Furthermore, we also analysed 13 molecular data from 13 major cancer types from The Cancer Genome Atlas (TGCA, *https://tcga-data.nci.nih.gov/tcga/*, date: 4/16/2016) project with survival information available. These cancer types include bladder urothelial carcinoma (BLCA), breast cancer carcinoma (BRCA), colon adenocarcinoma (COAD), head and neck squamous cell carcinoma (HNSC), kidney renal clear cell carcinoma (KIRC), acute myeloid leukemia (LAML), brain lower grade glioma (LGG), lung adenocarcinoma (LUAD), lung squamous cell carcinoma (LUSC), ovarian serous cystadenocarcinoma (OV), prostate adenocarcinoma (PRAD), skin cutaneous melanoma (SKCM), thyroid carcinoma (THCA), and uterine corpus endometrial carcinoma (UCEC). Each dataset contains 4 different types of molecular data, including protein expression, microRNA (miRNA) expression, mRNA expression (RNA-seq V2) and somatic copy number alterations (SCNAs). Note that the numbers of subjects varies in different data types.

## IV. ADDITIONAL NUMERICAL RESULTS

In this section, we provide more numerical results on ECC in terms of synthetic datasets evaluation, the number of basic partitions and the basic partition generation strategy.

### A. Performance of different clustering algorithms on synthetic data

Figure S3 and Figure S4 show the performance of different clustering algorithms on the 110 synthetic datasets with different settings. We find that for most datasets ECC achieves better performance than both traditional and previous ensemble clustering methods. For the four cases when ECC is not the best, the difference between the best one and ECC is within a small margin.

### B. Performance of different clustering algorithms on benchmark data sets

Table S V and Table SVI show the performance of different clustering algorithms in terms of R_n_ and *NMI*.

### C. Impact of the basic partition number

To fully uncover the properties of ECC for practical use, we thoroughly explore some impact factors of ECC. Fig. S5 shows the performance of ECC as a function of the number *r*. Generally speaking, the performance goes up with larger *r*, and the variance becomes smaller. The number of basic partitions determines the stability of ECC. We find that *r* = 100 is large enough for a robust partition.

### D. Comparison of different basic partition generation strategies

So far we employ Random Parameter Selection (RPS) strategy to generate basic partitions by *K*-means with different cluster numbers. A complementary strategy is the so-called Random Feature Selection (RFS) strategy. In RFS, we generate a subdata set by randomly selecting certain percentage of features (e.g. gene expressions), where *K*-means with a fixed cluster number is conducted to obtain the basic partitions. Fig. S6 demonstrates the performance of these two generation strategies. For these four datasets (15, 16, 26 and 28), RFS has better performance than RPS on all percentages of sampling ratio. For example, the improvements on dataset-15 (*Gordon-2002*) are over 80% and 40% in terms of *R_n_* and *NMI*. This indicates that only a subset of features (genes) reflect the true cluster structure, the rest are irrelevant or noisy. Finding discriminative features (genes) is a very challenging task, especially in unsupervised scenarios. Here we employ the simple RFS strategy and fuse these basic partitions to obtain promising results. Taking the efficiency into account, 10% sampling ratio is good enough for a satisfactory partition. Fig. S7 shows the comparison between the results derived from RPS and RFS (with 10% sampling ratio). We find that indeed RFS is a complementary basic partition generation strategy of RPS.

### E. Performance with missing values

We validate the performance of ECC in the presence of missing values, which result in incomplete basic partitions (IBPs). Given a dataset, we randomly remove certain instances and call*K*-means clustering algorithm on the rest instances with the user-defined cluster number. For these removed instances, the labels are assigned to be 0 in the incomplete basic partitions. We repeat the above process 100 times to obtain 100 IBPs and employ ECC to get the consensus one. Fig. S9 shows the performance of ECC with different missing ratios on 4 datesets. We find that ECC can still provide high quality and robust consensus partition even with high missing ratio.

## V. SURVIVAL ANALYSIS

For real-world molecular data without label information (e.g. the 13 TCGA cancer types analyzed in this work), we can employ survival analyses to evaluate the performance of different clustering methods. Survival analysis considers the expected duration of time until one or more events happen, such as death, disease occurrence, disease recurrence, recovery, or other experience of interest[5]. The duration of time measures the time from the beginning of an observation period (such as surgery or beginning treatment) to an event, or end of the study, or loss of contact or withdrawal from the study. Censoring/Censored observation means that, if a subject does not have an event during the observation time, they are described as censored. The subject is censored in the sense that nothing is observed or known about that subject after the time of censoring. A censored subject may or may not have an event after the end of observation time.

### A. Log-rank test

The log-rank test is a hypothesis test to compare the survival distributions of two or more groups. The null hypothesis that every group has the same survival function. The expected number of subjects surviving at each time point in each group is adjusted for the number of subjects at risk in the groups at each event time. The log-rank test determines if the observed number of events in each group is significantly different from the expected number. The formal test is based on a chi-squared statistic. The log-rank statistic has a chi-squared distribution with one degree of freedom, and the p-value is calculated using the chi-squared distribution. When the p-value is smaller than 0.05, it typically indicates that those groups differ significantly in survival times.

### B. Survival analysis on real-world data

Tables S7-10 display the log-rank p-values for survival analysis of 13 TCGA major cancer types using different molecular data type: mRNA expression, microRNA expression, protein expression, and somatic copy number alterations (SCNAs); and different clustering methods. The p-values that are smaller than 0.05 are displayed in bold face. We found that for each single molecular data type our ECC method yields more significant p-values than other clustering methods. Table S11 displays the log-rank p-values for survival analysis of 13 TCGA major cancer types using pan-omics data (i.e. integrating mRNA expression, microRNA expression, protein expression, and SCNAs) and our ECC method. (Note that the competitive clustering methods cannot handle missing values or incomplete basic partitions, hence we only show the result of ECC on the pan-omics data.) We find that for each cancer type the p-value is smaller than 0.05, suggesting that by integrating 4 different molecular data types ECC generated significant subtypes for all the 13 cancer types.

---

**Table SI.** Notations for Supplementary Note 1.

| Symbol | Domain | Description |
| --- | --- | --- |
| $n$ | $\mathbf{Z}$ | the number of subjects |
| $m$ | $\mathbf{Z}$ | the number of genes |
| $r$ | $\mathbf{Z}$ | the number of basic partitions |
| $\mathcal{X}$ | $\mathbf{R}^{n \times m}$ | the gene expression data matrix |
| $\pi_v$ | $\{1, 2, \dots, K\}^n$ | the $v$-th basic partition |
| $K_v$ | $\mathbf{Z}$ | the cluster number of $\pi_v$ |
| $\pi$ | $\{1, 2, \dots, K\}^n$ | the consensus partition |
| $K$ | $\mathbf{Z}$ | the cluster number of $\pi$ |
| $B$ | $0, 1^{n \times \sum_{v=1}^r K_v}$ | the binary matrix for clustering |
| $m_k$ | $\mathbf{R}^{\sum_{v=1}^r K_v}$ | the $k$-th centroid matrix |

**Table SII.** Notations for Supplementary Note 2.

| Symbol | Domain | Description |
| --- | --- | --- |
| $n$ | $\mathbf{Z}$ | the number of genes in the network |
| $F_i^{\text{mRNA}}$ | $\mathbf{R}$ | the changing rate of mRNA concentration |
| $F_i^{\text{Prot}}$ | $\mathbf{R}$ | the changing rate of protein concentration |
| $m_i$ | $[0,1]$ | the maximum transcription rate |
| $r_i$ | $[0,1]$ | the translation rate |
| $\lambda_i^{\text{mRNA}}$ | $\mathbf{R}$ | mRNA degradation rate |
| $\lambda_i^{\text{Prot}}$ | $\mathbf{R}$ | protein degradation rate |
| $f_i(\cdot)$ | $[0,1]$ | the activation function of gene $i$ |
| $k_{ij}$ | $\mathbf{R}$ | the dissociation constant |
| $h_{ij}$ | $[1,10]$ | the Hill coefficient |
| $\alpha_i$ | $\mathbf{R}$ | the relative activation in state $S_0$ |
| $\rho$ | $\mathbf{R}$ | the cooperativity factor |
| $\eta_v, \eta_d, \eta_{\omega_i}$ | $\mathbf{R}$ | independent Gaussian white-noise process |
| $c_1$ | $(0,1)$ | molecular noise level |
| $c_2$ | $(0,1)$ | measurement noise level |
| $N$ | $\mathbf{Z}$ | the number of samples |
| $p$ | $\mathbf{Z}$ | the number of knock-out genes |

**Table SIII.** Some key characteristics of 35 benchmark datasets for cluster validity.

| No | Dataset | Tissue | #Subject | #Genes | #Class | No | Dataset | Tissue | #Subject | #Genes | #Class |
| --- | --- | --- | --- | --- | --- | --- | --- | --- | --- | --- | --- |
| 1 | Alizadeh-2000-v1 | Blood | 42 | 1095 | 2 | 19 | Lapointe-2004-v2 | Prostate | 110 | 2496 | 4 |
| 2 | Alizadeh-2000-v2 | Blood | 62 | 2093 | 3 | 20 | Liang-2005 | Brain | 37 | 1411 | 3 |
| 3 | Alizadeh-2000-v3 | Blood | 62 | 2093 | 4 | 21 | Nutt-2003-v1 | Brain | 50 | 1377 | 4 |
| 4 | Armstrong-2002-v1 | Blood | 72 | 1081 | 2 | 22 | Nutt-2003-v2 | Brain | 28 | 1070 | 2 |
| 5 | Armstrong-2002-v2 | Blood | 72 | 2194 | 3 | 23 | Nutt-2003-v3 | Brain | 22 | 1152 | 2 |
| 6 | Bhattacharjee-2001 | Lung | 203 | 1543 | 5 | 24 | Pomeroy-2002-v1 | Brain | 34 | 857 | 2 |
| 7 | Bitter-2000 | Skin | 38 | 2201 | 2 | 25 | Pomeroy-2002-v2 | Brain | 42 | 1379 | 5 |
| 8 | Bredel-2005 | Brain | 50 | 1739 | 3 | 26 | Ramaswamy-2001 | Multi-tissue | 190 | 1363 | 14 |
| 9 | Chen-2002 | Liver | 179 | 85 | 2 | 27 | Risinger-2003 | Endometrium | 42 | 1771 | 4 |
| 10 | Chowdary-2006 | Multi-tissue | 104 | 182 | 2 | 28 | Shipp-2002 | Blood | 77 | 798 | 2 |
| 11 | Dyrskjot-2003 | Bladder | 40 | 1203 | 3 | 29 | Singh-2002 | Prostate | 102 | 339 | 2 |
| 12 | Garber-2001 | Lung | 66 | 4553 | 4 | 30 | Su-2001 | Multi-tissue | 174 | 1571 | 10 |
| 13 | Golub-1999-v1 | Bone marrow | 72 | 1868 | 2 | 31 | Tomlins-2006-v1 | Prostate | 92 | 1288 | 4 |
| 14 | Golub-1999-v2 | Bone marrow | 72 | 1868 | 3 | 32 | Tomlins-2006-v2 | Prostate | 104 | 2315 | 5 |
| 15 | Gordon-2002 | Lung | 181 | 1626 | 2 | 33 | West-2001 | Breast | 49 | 1198 | 2 |
| 16 | Khan-2001 | Multi-tissue | 83 | 1069 | 4 | 34 | Yeoh-2002-v1 | Bone marrow | 248 | 2526 | 2 |
| 17 | Laiho-2007 | Colon | 37 | 2202 | 2 | 35 | Yeoh-2002-v2 | Bone marrow | 248 | 2526 | 6 |
| 18 | Lapointe-2004-v1 | Prostate | 69 | 1625 | 3 |  |  |  |  |  |  |

**Table SIV.** Some key characteristics of 13 real-world datasets from TCGA. Note: the cluster number is obtain from the original papers which publish the datasets.

| No | Database | #Class | protein |  | miRNA |  | mRNA |  | SCNA |  |
| --- | --- | --- | --- | --- | --- | --- | --- | --- | --- | --- |
|  |  |  | #Instance | #protein | #Instance | #miRNA | #Instance | #mRNA | #Instance | #SCNA |
| 1 | BLCA [6] | 4 | 127 | 190 | 328 | 1046 | 326 | 20531 | 330 | 24952 |
| 2 | BRCA [7] | 4 | 742 | 190 | 728 | 1046 | 1065 | 20531 | 1067 | 24952 |
| 3 | COAD [8] | 4 | 330 | 190 | 242 | 1046 | 292 | 20531 | 450 | 24952 |
| 4 | HNSC [9] | 4 | 212 | 190 | 471 | 1046 | 502 | 20531 | 508 | 24952 |
| 5 | KIRC [10] | 4 | 454 | 190 | 247 | 1046 | 523 | 20531 | 524 | 24952 |
| 6 | LGG [11] | 3 | 258 | 190 | 441 | 1046 | 445 | 20531 | 443 | 24952 |
| 7 | LUAD [12] | 3 | 237 | 190 | 441 | 1046 | 496 | 20531 | 502 | 24952 |
| 8 | LUSC [13] | 4 | 195 | 190 | 317 | 1046 | 476 | 20531 | 479 | 24952 |
| 8 | OV [14] | 4 | 408 | 190 | 474 | 1046 | 262 | 20531 | 575 | 24952 |
| 10 | PRAD [15] | 7 | 161 | 190 | 414 | 1046 | 418 | 20531 | 418 | 24952 |
| 11 | SCKM [16] | 4 | 206 | 190 | 416 | 1046 | 436 | 20531 | 438 | 24952 |
| 12 | THCA [17] | 5 | 370 | 190 | 503 | 1046 | 502 | 20531 | 503 | 24952 |
| 13 | UCEC [18] | 4 | 394 | 190 | 393 | 1046 | 162 | 20531 | 527 | 24952 |

**Table SV.** Performance of di.erent clustering algorithms on benchmark datasets by *R_n_*

| No | Dataset | AL | SL | CL | KM | SC | LCE | ASRS | ECC |
| --- | --- | --- | --- | --- | --- | --- | --- | --- | --- |
| 1 | Alizadeh-2000-v1 | 0.0047 | 0.0000 | 0.0047 | 0.1856 | 0.1633 | 0.4981 | 0.4306 | **0.5576** |
| 2 | Alizadeh-2000-v2 | **0.8519** | -0.0427 | 0.7483 | 0.5241 | 0.5059 | 0.4100 | 0.4091 | 0.4093 |
| 3 | Alizadeh-2000-v3 | 0.4387 | -0.0129 | 0.3848 | 0.3917 | 0.3892 | 0.3235 | **0.4615** | 0.4464 |
| 4 | Armstrong-2002-v1 | -0.0451 | -0.0138 | **0.4753** | 0.2356 | 0.2680 | 0.2102 | 0.2384 | 0.2296 |
| 5 | Armstrong-2002-v2 | 0.0009 | 0.0007 | 0.4620 | 0.5690 | 0.5876 | 0.7821 | 0.7747 | **0.8480** |
| 6 | Bhattacharjee-2001 | 0.2606 | 0.0390 | **0.4662** | 0.2173 | 0.1962 | 0.2040 | 0.2441 | 0.2362 |
| 7 | Bitter-2000 | 0.0000 | 0.0000 | -0.0103 | -0.0144 | -0.0157 | 0.0179 | 0.0179 | **0.0429** |
| 8 | Bredel-2005 | 0.0056 | 0.0056 | 0.3671 | 0.3606 | 0.2933 | 0.3270 | **0.5183** | 0.3766 |
| 9 | Chen-2002 | -0.0031 | -0.0031 | -0.0065 | 0.2571 | -0.0065 | 0.5365 | -0.006 | **0.6505** |
| 10 | Chowdary-2006 | 0.0091 | 0.0091 | 0.0091 | 0.0657 | 0.0091 | 0.5273 | 0.0657 | **0.8857** |
| 11 | Dyrskjot-2003 | 0.0534 | 0.0534 | 0.0534 | 0.5069 | 0.5563 | 0.5779 | 0.5077 | **0.6213** |
| 12 | Garber-2001 | -0.0161 | -0.0161 | 0.1023 | 0.1766 | **0.1776** | 0.0722 | 0.1558 | 0.1084 |
| 13 | Golub-1999-v1 | 0.2313 | 0.0243 | 0.5016 | 0.5646 | 0.6860 | 0.6347 | 0.5905 | **0.8473** |
| 14 | Golub-1999-v2 | 0.1710 | 0.0138 | 0.2440 | 0.5456 | 0.5758 | 0.6709 | **0.8776** | 0.8127 |
| 15 | Gordon-2002 | -0.0086 | -0.0086 | -0.0882 | -0.0618 | -0.0168 | -0.0583 | -0.0948 | **-0.0126** |
| 16 | Khan-2001 | -0.0126 | -0.0152 | 0.1848 | 0.2979 | 0.5388 | 0.2951 | 0.3394 | **0.4433** |
| 17 | Laiho-2007 | 0.0838 | 0.1355 | -0.1028 | **0.1964** | 0.1914 | 0.1914 | 0.1244 | 0.1284 |
| 18 | Lapointe-2004-v1 | 0.0402 | 0.0402 | 0.0402 | **0.1770** | 0.1549 | 0.1037 | 0.1568 | 0.1295 |
| 19 | Lapointe-2004-v2 | 0.0253 | 0.0130 | 0.0375 | 0.1196 | 0.0798 | 0.0968 | 0.1018 | **0.1386** |
| 20 | Liang-2005 | 0.0497 | -0.1529 | 0.1573 | 0.1551 | 0.1775 | **0.1937** | 0.1573 | 0.1573 |
| 21 | Nutt-2003-v1 | 0.0008 | 0.0005 | 0.1232 | 0.3246 | 0.2956 | 0.2483 | 0.3562 | **0.3766** |
| 22 | Nutt-2003-v2 | 0.0023 | 0.0000 | 0.1082 | 0.0314 | 0.0023 | 0.3055 | 0.0023 | **0.4176** |
| 23 | Nutt-2003-v3 | 0.1020 | 0.1020 | 0.1020 | 0.6993 | 0.8153 | 0.8153 | **1.0000** | 0.8153 |
| 24 | Pomeroy-2002-v1 | -0.0645 | **-0.0366** | -0.0459 | -0.0513 | -0.0459 | -0.0459 | -0.0459 | -0.0468 |
| 25 | Pomeroy-2002-v2 | 0.0975 | 0.1012 | 0.2549 | 0.4636 | 0.5353 | 0.4571 | 0.4748 | **0.4979** |
| 26 | Ramaswamy-2001 | -0.0021 | -0.0029 | 0.0126 | 0.1241 | 0.2221 | 0.1782 | 0.1348 | **0.2285** |
| 27 | Risinger-2003 | -0.0272 | -0.0108 | -0.0640 | 0.1122 | 0.1088 | 0.1000 | 0.0989 | **0.1212** |
| 28 | Shipp-2002 | **-0.0325** | **-0.0325** | **-0.0325** | -0.0848 | **-0.0325** | -0.0935 | -0.1003 | -0.0918 |
| 29 | Singh-2002 | **0.0269** | 0.0008 | **0.0269** | 0.0259 | 0.0330 | 0.0047 | 0.0245 | 0.0044 |
| 30 | Su-2001 | 0.1296 | 0.0337 | 0.1821 | 0.3831 | 0.4959 | 0.4007 | 0.4175 | **0.4532** |
| 31 | Tomlins-2006-v1 | 0.0180 | 0.0122 | 0.0485 | 0.1643 | 0.1747 | **0.1911** | 0.1510 | 0.1305 |
| 32 | Tomlins-2006-v2 | 0.0099 | 0.0099 | 0.3178 | 0.2845 | 0.2339 | 0.3369 | **0.4154** | 0.3726 |
| 33 | West-2001 | 0.0001 | -0.0016 | 0.0001 | 0.2834 | 0.0001 | **0.3875** | **0.3875** | **0.3875** |
| 34 | Yeoh-2002-v1 | -0.0063 | -0.0063 | -0.0063 | 0.3664 | 0.0846 | 0.8803 | -0.0063 | **0.9344** |
| 35 | Yeoh-2002-v2 | -0.0011 | -0.0011 | -0.0011 | 0.1785 | 0.1933 | 0.1277 | 0.1866 | **0.2376** |
| Average |  | 0.0684 | 0.0068 | 0.1445 | 0.2507 | 0.2465 | 0.3117 | 0.2734 | **0.3684** |

**Table SVI.** Performance of di.erent clustering algorithms on benchmark datasets by *R_n_*

| No | Dataset | AL | SL | CL | KM | SC | LCE | ASRS | ECC |
| --- | --- | --- | --- | --- | --- | --- | --- | --- | --- |
| 1 | Alizadeh-2000-v1 | 0.0939 | 0.0601 | 0.0939 | 0.1776 | 0.1369 | 0.4167 | 0.3516 | **0.4940** |
| 2 | Alizadeh-2000-v2 | **0.8119** | 0.0342 | 0.6972 | 0.6454 | 0.6634 | 0.5672 | 0.5877 | 0.5832 |
| 3 | Alizadeh-2000-v3 | 0.6421 | 0.0960 | 0.5665 | 0.5789 | 0.5930 | 0.5300 | **0.6441** | 0.6340 |
| 4 | Armstrong-2002-v1 | 0.0631 | 0.0263 | **0.4389** | 0.3606 | 0.3793 | 0.3459 | 0.3622 | 0.3571 |
| 5 | Armstrong-2002-v2 | 0.1311 | 0.0780 | 0.5536 | 0.6133 | 0.6243 | 0.7654 | 0.7460 | **0.8154** |
| 6 | Bhattacharjee-2001 | 0.4039 | 0.1014 | **0.5032** | 0.4252 | 0.4504 | 0.4186 | 0.4941 | 0.4585 |
| 7 | Bitter-2000 | **0.0640** | **0.0640** | 0.0000 | 0.0127 | 0.0083 | 0.0323 | 0.0323 | 0.0502 |
| 8 | Bredel-2005 | 0.0848 | 0.0848 | 0.3685 | **0.3809** | 0.2916 | 0.3688 | 0.4769 | 0.3712 |
| 9 | Chen-2002 | 0.0199 | 0.0199 | 0.0008 | 0.2058 | 0.0008 | 0.4980 | 0.0010 | **0.6273** |
| 10 | Chowdary-2006 | 0.0459 | 0.0459 | 0.0459 | 0.1424 | 0.0459 | 0.4739 | 0.1424 | **0.8098** |
| 11 | Dyrskjot-2003 | 0.1613 | 0.1613 | 0.1613 | 0.5124 | 0.5082 | **0.5743** | 0.5158 | 0.5584 |
| 12 | Garber-2001 | 0.0734 | 0.0734 | 0.0999 | 0.1801 | 0.1689 | 0.1647 | 0.1294 | **0.2025** |
| 13 | Golub-1999-v1 | 0.2764 | 0.0684 | 0.4837 | 0.5358 | 0.5809 | 0.6085 | 0.5536 | **0.8149** |
| 14 | Golub-1999-v2 | 0.2868 | 0.0803 | 0.3496 | 0.5762 | 0.5429 | 0.6870 | **0.8453** | 0.7737 |
| 15 | Gordon-2002 | 0.0083 | 0.0083 | 0.1168 | 0.1385 | **0.1684** | 0.1411 | 0.1099 | 0.1479 |
| 16 | Khan-2001 | 0.0966 | 0.0941 | 0.4857 | 0.5149 | **0.6718** | 0.5109 | 0.5644 | 0.6221 |
| 17 | Laiho-2007 | 0.0340 | 0.1680 | 0.0664 | 0.1479 | **0.1959** | **0.1959** | 0.0749 | 0.0768 |
| 18 | Lapointe-2004-v1 | 0.0466 | 0.0466 | 0.0466 | **0.1956** | 0.1269 | 0.1101 | 0.1826 | 0.1281 |
| 19 | Lapointe-2004-v2 | 0.0513 | 0.0406 | 0.0419 | 0.1620 | 0.1173 | 0.1357 | 0.1379 | **0.1652** |
| 20 | Liang-2005 | 0.3002 | 0.1181 | 0.3545 | **0.3811** | 0.3597 | 0.3772 | 0.3545 | 0.3545 |
| 21 | Nutt-2003-v1 | 0.1293 | 0.1296 | 0.2900 | 0.4742 | 0.4233 | 0.3844 | **0.5083** | 0.4994 |
| 22 | Nutt-2003-v2 | 0.0406 | 0.0778 | 0.2513 | 0.0589 | 0.0406 | 0.3988 | 0.0406 | **0.4021** |
| 23 | Nutt-2003-v3 | 0.1600 | 0.1600 | 0.1600 | 0.6545 | 0.7523 | 0.7523 | **1.0000** | 0.7523 |
| 24 | Pomeroy-2002-v1 | 0.0521 | 0.0332 | 0.0079 | 0.0453 | 0.0079 | 0.0079 | 0.0079 | **0.0092** |
| 25 | Pomeroy-2002-v2 | 0.4150 | 0.4177 | 0.5396 | 0.5935 | 0.6401 | 0.6032 | 0.5937 | **0.6171** |
| 26 | Ramaswamy-2001 | 0.2176 | 0.1608 | 0.3376 | 0.5203 | 0.5172 | 0.5126 | **0.5560** | 0.5277 |
| 27 | Risinger-2003 | 0.1349 | 0.1203 | 0.1877 | 0.2867 | 0.2744 | 0.2784 | 0.2709 | **0.2911** |
| 28 | Shipp-2002 | 0.0288 | 0.0288 | 0.0288 | 0.0747 | 0.0288 | 0.1235 | 0.1124 | **0.1280** |
| 29 | Singh-2002 | 0.0675 | 0.0360 | **0.0675** | 0.0475 | 0.0562 | 0.0101 | 0.0342 | 0.0100 |
| 30 | Su-2001 | 0.4919 | 0.2874 | 0.5088 | 0.6018 | **0.6614** | 0.6122 | 0.6422 | 0.6276 |
| 31 | Tomlins-2006-v1 | 0.1281 | 0.1058 | 0.1574 | 0.2849 | 0.2533 | **0.3257** | 0.3065 | 0.2558 |
| 32 | Tomlins-2006-v2 | 0.1158 | 0.1158 | 0.4676 | 0.4653 | 0.3586 | 0.5050 | **0.5815** | 0.5266 |
| 33 | West-2001 | 0.0823 | 0.0530 | 0.0823 | 0.2603 | 0.0823 | **0.3125** | **0.3125** | **0.3125** |
| 34 | Yeoh-2002-v1 | 0.0070 | 0.0070 | 0.0070 | 0.3342 | 0.0366 | 0.7968 | 0.0070 | **0.8564** |
| 35 | Yeoh-2002-v2 | 0.0691 | 0.0691 | 0.0691 | 0.3760 | 0.3309 | 0.2768 | **0.4175** | 0.3815 |
| Average |  | 0.1667 | 0.0935 | 0.2468 | 0.3419 | 0.3171 | 0.3949 | 0.3628 | **0.4355** |

**Table SVII.**
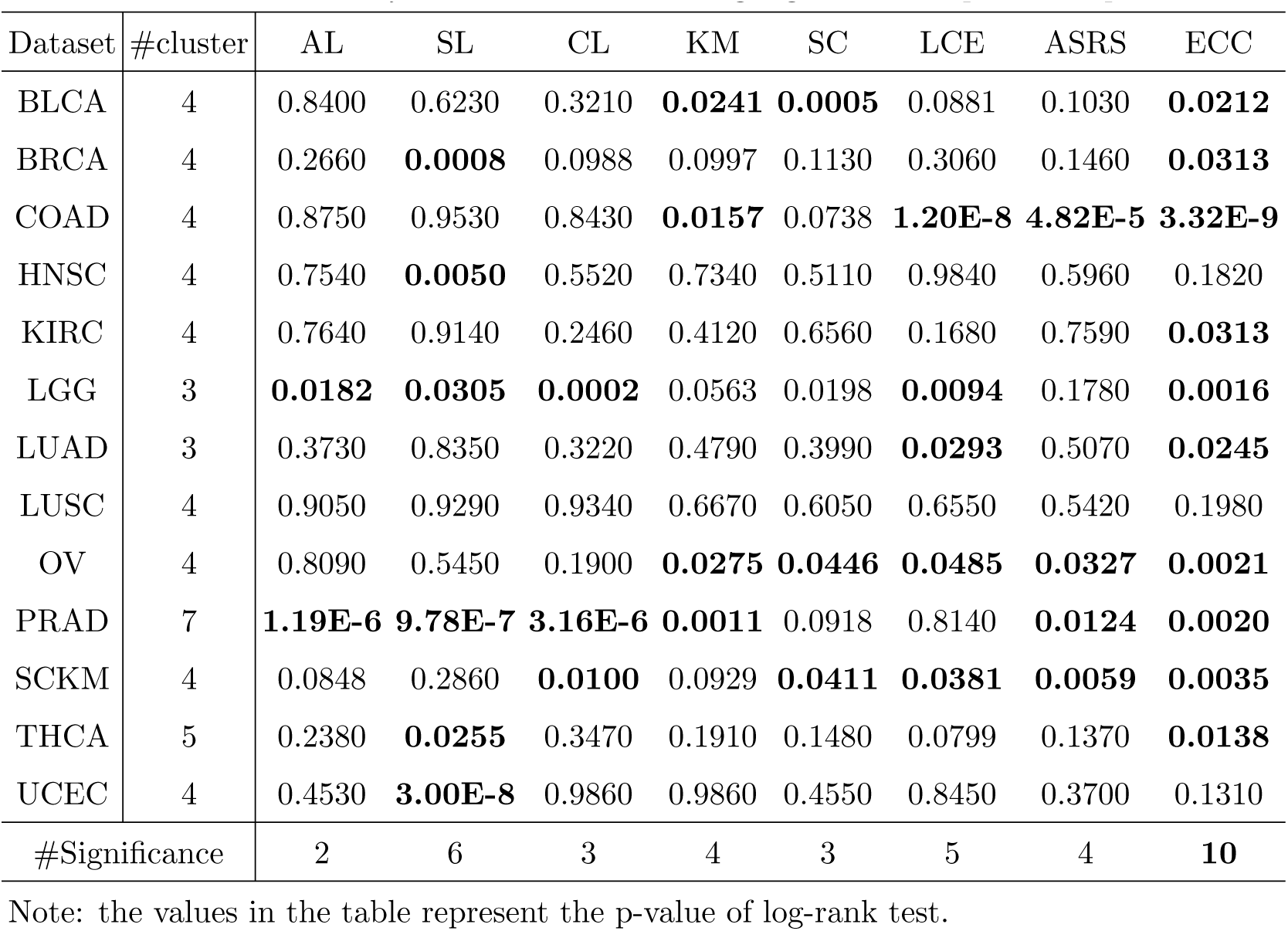
Survival analysis of different clustering algorithms on protein expression data. Note: the values in the table represent the p-value of log-rank test.

**Table SVIII.** Survival analysis of di.erent clustering algorithms on miRNA expression data. Note: the values in the table represent the p-value of log-rank test.

| Dataset | #cluster | AL | SL | CL | KM | SC | LCE | ASRS | ECC |
| --- | --- | --- | --- | --- | --- | --- | --- | --- | --- |
| BLCA | 4 | 0.2780 | 0.5880 | 0.5940 | 0.0616 | 0.5620 | 0.3410 | 0.2400 | **0.0124** |
| BRCA | 4 | 0.3110 | 0.6350 | 0.5410 | **1.53E-5** | 0.0717 | **3.97E-6** | **1.12E-7** | **6.37E-8** |
| COAD | 4 | 0.3290 | 0.6430 | 0.2070 | 0.2290 | 0.1960 | **8.88E-4** | **0.0246** | **5.91E-4** |
| HNSC | 4 | 0.8900 | 0.8820 | 0.7650 | 0.5760 | 0.6770 | 0.0605 | **4.45E-5** | **0.0090** |
| KIRC | 4 | 0.7970 | 0.6420 | 0.0692 | 0.2180 | **0.0093** | **0.0180** | 0.1090 | **0.0223** |
| LGG | 3 | 0.8820 | 0.9640 | 0.8940 | 0.9850 | 0.9000 | 0.7450 | 0.0640 | 0.0751 |
| LUAD | 3 | 0.8350 | 0.1200 | 0.7410 | 0.2870 | 0.3580 | **0.0038** | 0.8260 | **0.0028** |
| LUSC | 4 | 0.1060 | 0.3450 | 0.0565 | **0.0152** | **0.0394** | 0.1310 | 0.3120 | **0.0442** |
| OV | 4 | 0.5540 | **0.0007** | 0.2410 | 0.6290 | 0.4190 | 0.2340 | 0.2340 | **0.0375** |
| PRAD | 7 | 0.4570 | 0.4250 | 0.6500 | 0.3330 | 0.3200 | 0.8720 | 0.6270 | 0.1840 |
| SCKM | 4 | 0.0619 | 0.6870 | 0.4920 | 0.6390 | 0.6940 | 0.0663 | 0.0575 | **0.0076** |
| THCA | 5 | 0.4660 | **0.0064** | **0.0053** | 0.0892 | 0.1100 | **0.0119** | **0.0157** | **3.75E-4** |
| UCEC | 4 | 0.5280 | 0.4570 | 0.6290 | 0.6870 | 0.6080 | 0.5530 | 0.3520 | 0.2680 |
| #Significance |  | 0 | 2 | 1 | 2 | 2 | 5 | 4 | **10** |

Note: the values in the table represent the p-value of log-rank test.

**Table SIX.** Survival analysis of di.erent clustering algorithms on mRNA expression data. Note: the values in the table represent the p-value of log-rank test.

| Dataset | #cluster | AL | SL | CL | KM | SC | LCE | ASRS | ECC |
| --- | --- | --- | --- | --- | --- | --- | --- | --- | --- |
| BLCA | 4 | **1.06E-7** | **8.88E-8** | **1.06E-7** | **0.0258** | 0.6860 | 0.1280 | 0.0938 | **0.0187** |
| BRCA | 4 | **5.35E-3** | 0.1740 | **0.0401** | 0.1760 | 0.0840 | 0.5980 | **0.0155** | **0.0011** |
| COAD | 4 | 0.8930 | 0.8960 | 0.8720 | **0.0163** | **0.0296** | **0.0048** | 0.0743 | **0.0008** |
| HNSC | 4 | 0.2950 | **8.53E-5** | 0.1350 | 0.7470 | 0.5440 | 0.6290 | 0.1440 | 0.1160 |
| KIRC | 4 | **0.0025** | **0.0012** | **0.0036** | 0.0612 | 0.1450 | 0.2420 | 0.1550 | **0.0314** |
| LGG | 3 | **0.0156** | **0.0156** | **0.0155** | 0.1270 | 0.1230 | 0.2650 | **0.0023** | **0.0039** |
| LUAD | 3 | **0.0109** | 0.8290 | 0.3190 | **0.0429** | **0.0034** | **0.0189** | **0.0157** | **0.0028** |
| LUSC | 4 | 0.0990 | 0.2100 | **0.0241** | **0.0355** | 0.4740 | 0.0769 | 0.1360 | **0.0258** |
| OV | 4 | 0.2210 | **4.92E-10** | 0.1700 | 0.6360 | 0.3780 | 0.8720 | 0.7660 | 0.2210 |
| PRAD | 7 | **4.29E-9** | **4.49E-9** | **5.88E-9** | **7.29E-11** | **4.10E-9** | **0.0070** | **6.75E-13** | **8.59E-6** |
| SCKM | 4 | **0.0012** | **0.0012** | **0.0015** | 0.5230 | **0.0006** | 0.1350 | **5.91E-10** | **3.94E-6** |
| THCA | 5 | **0.0147** | 0.5650 | 0.0713 | **0.0244** | 0.0561 | 0.2380 | 0.0710 | **0.0024** |
| UCEC | 4 | 0.5790 | 0.0594 | 0.1930 | 0.1850 | 0.2460 | 0.3670 | 0.4890 | 0.1240 |
| #Significance |  | 8 | 7 | 7 | 6 | 4 | 3 | 5 | **10** |

Note: the values in the table represent the p-value of log-rank test.

**Table SX.** Survival analysis of di.erent clustering algorithms on SCNA data. Note: the values in the table represent the p-value of log-rank test.

| Dataset | #cluster | AL | SL | CL | KM | SC | LCE | ASRS | ECC |
| --- | --- | --- | --- | --- | --- | --- | --- | --- | --- |
| BLCA | 4 | 0.3710 | 0.3710 | 0.3810 | 0.6340 | 0.3580 | 0.4340 | 0.3800 | 0.1910 |
| BRCA | 4 | 0.6540 | 0.6540 | 0.1160 | **0.0090** | 0.4790 | 0.0798 | 0.3520 | **0.0375** |
| COAD | 4 | 0.9320 | 0.9320 | 0.9010 | 0.1600 | 0.7920 | 0.7670 | 0.4660 | 0.2340 |
| HNSC | 4 | **0.0003** | **0.0003** | **0.0380** | 0.5280 | 0.5730 | 0.8280 | 0.7710 | 0.3800 |
| KIRC | 4 | 0.6580 | 0.7510 | 0.0929 | 0.4390 | 0.1060 | 0.2690 | 0.3710 | 0.1730 |
| LGG | 3 | 0.8800 | 0.9950 | 0.6430 | 0.5710 | 0.6130 | 0.8750 | 0.9740 | 0.4130 |
| LUAD | 3 | 0.5420 | 0.5420 | 0.5880 | 0.0763 | 0.2390 | **0.0121** | **0.0080** | **0.0067** |
| LUSC | 4 | 0.8900 | 0.8190 | 0.3870 | 0.3560 | 0.3810 | 0.1710 | 0.5540 | **0.0425** |
| OV | 4 | 0.7500 | 0.7500 | 0.1270 | 0.1710 | 0.0904 | 0.1730 | 0.1380 | **0.0359** |
| PRAD | 7 | 0.8410 | **2.40E-7** | 0.5060 | 0.2640 | **0.0008** | **0.0160** | **0.0046** | **0.0007** |
| SCKM | 4 | 0.8730 | 0.8140 | 0.6790 | 0.5660 | 0.1970 | 0.2210 | 0.2040 | **0.0491** |
| THCA | 5 | 0.1530 | 0.5180 | 0.1440 | 0.2670 | 0.1960 | 0.1360 | 0.5440 | 0.1080 |
| UCEC | 4 | 0.1100 | 0.1100 | 0.2310 | **0.0484** | 0.0673 | 0.4860 | 0.3450 | 0.1260 |
| #Significance |  | 1 | 2 | 1 | 2 | 1 | 2 | 2 | **6** |

Note: the values in the table represent the p-value of log-rank test.

**Table SXI.** Survival analysis of ECC on pan-omics data. Note: the values in the table represent the p-value of log-rank test.

| Dataset | #cluster | ECC | Dataset | #cluster | ECC | Dataset | #cluster | ECC |
| --- | --- | --- | --- | --- | --- | --- | --- | --- |
| BLCA | 4 | **0.0027** | BRCA | 4 | **0.0131** | COAD | 4 | **8.69E-6** |
| HNSC | 4 | **0.0323** | KIRC | 4 | **0.0010** | LGG | 3 | **0.0119** |
| LUAD | 3 | **2.87E-5** | LUSC | 4 | **0.0393** | OV | 4 | **0.0008** |
| PRAD | 7 | **0.0006** | SCKM | 4 | **0.0131** | THCA | 5 | **0.0035** |
| UCEC | 4 | **0.0043** |  |  |  |  |  |  |

Note: the values in the table represent the p-value of log-rank test.

**Figure S1.**
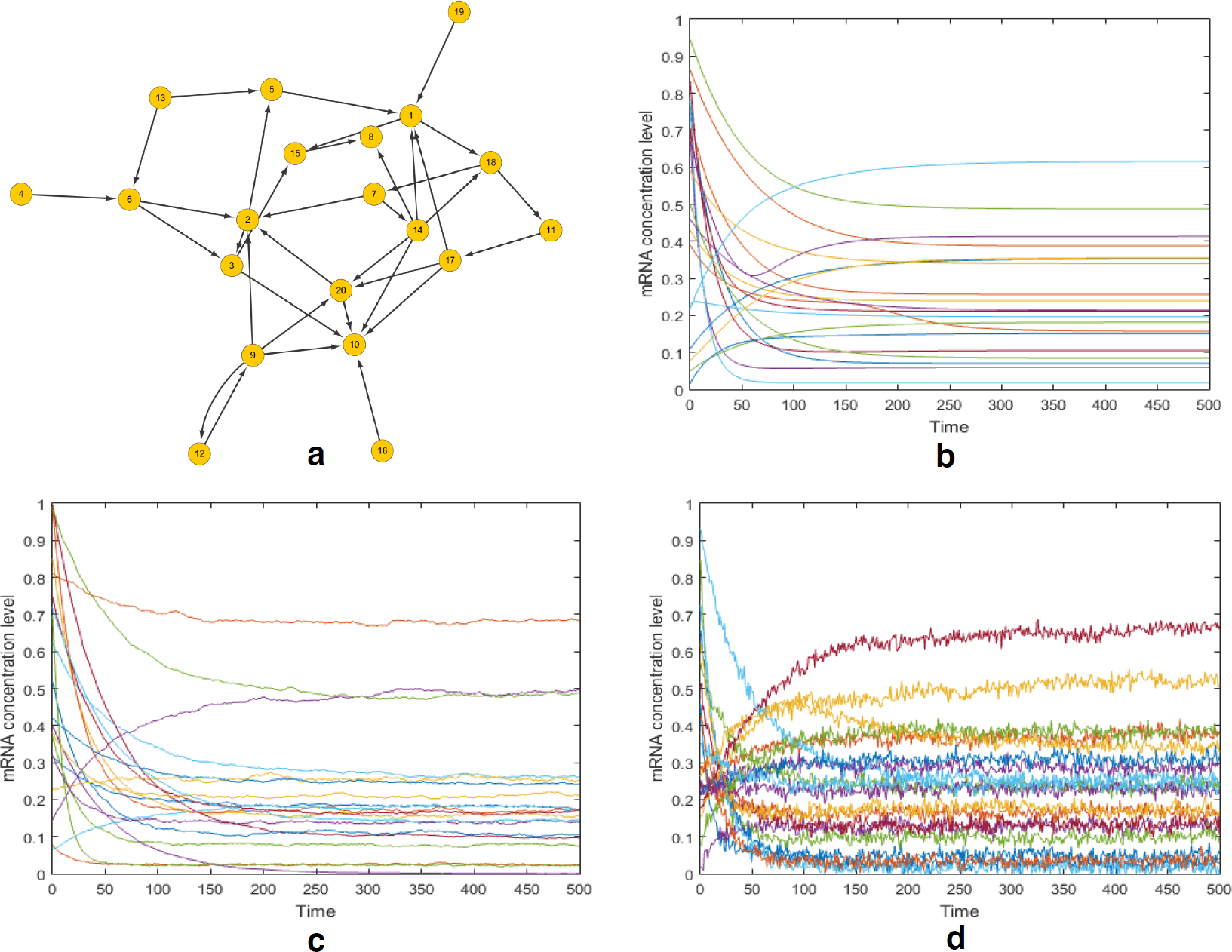
The random regulatory network of 20 genes and the time evolutions of mRNA concentration level. **a** represents the gene regulatory network of 20 genes, which is generated randomly with the mean in-degree of 2. **b** depicts the time evolution of mRNA concentration level without noise. The initial states are randomly generated. Each curve represents the time evolution of mRNA concentration level of a certain gene node. **c** depicts the time evolution of mRNA concentration level with molecular noise level = 0.01. **d** depicts the time evolution of mRNA concentration level with molecular noise level = 0.01, measurement noise level = 0.012. Each line represents the time evolution of mRNA concentration level of a gene node. It is easy to find that the time evolution curve without noise is smooth, while the time evolution with noise as a stochastic process is rough.

**Figure S2.**
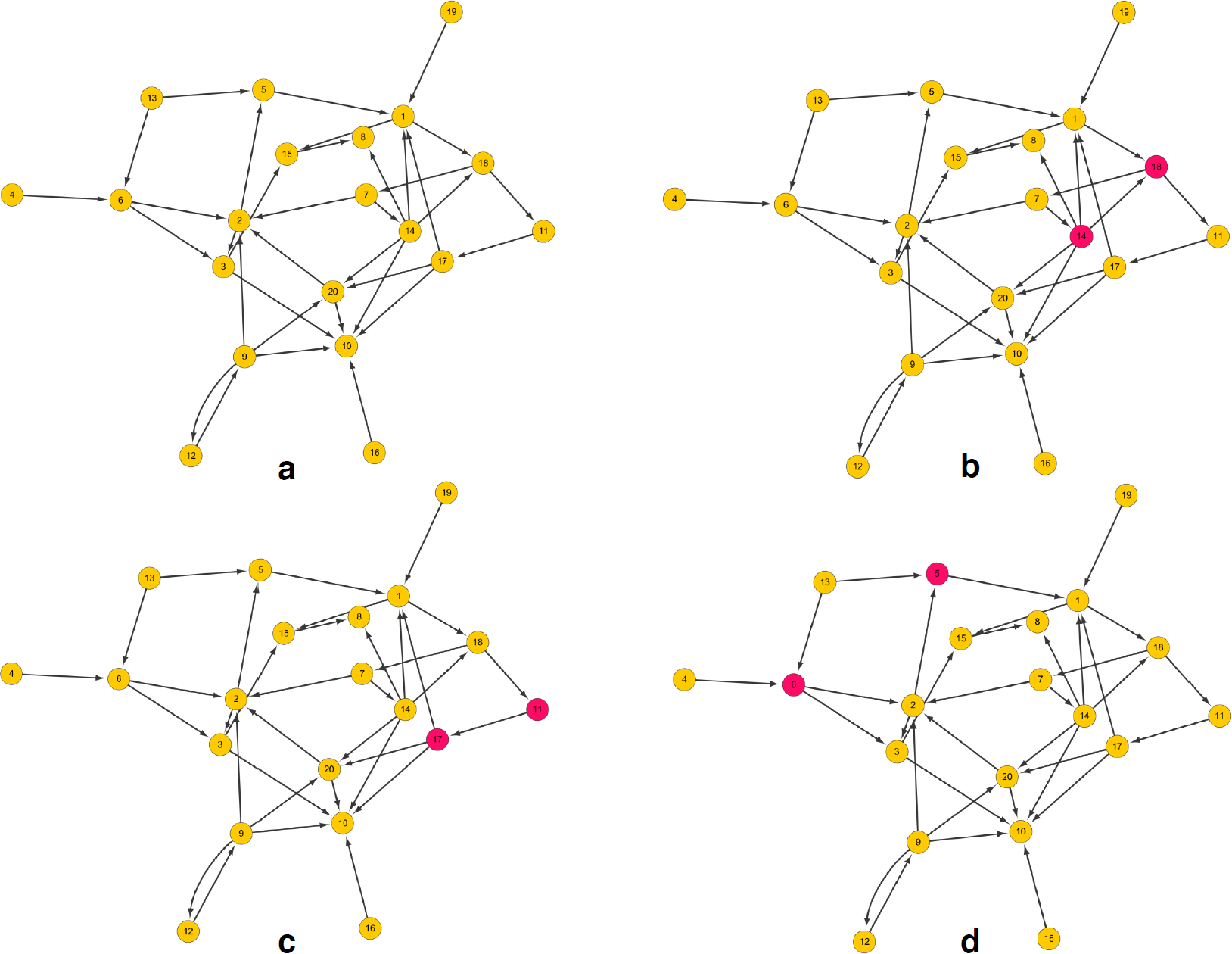
Normal gene regulatory network and three subtypes. **a** illustrates the normal gene regulatory network of 20 genes. **b** illustrates subtype 1 with two randomly knock-out genes. **c** illustrates subtype 2 with two randomly knock-out genes. **d** illustrates subtype 3 with two randomly knock-out genes.

**Figure S3.**
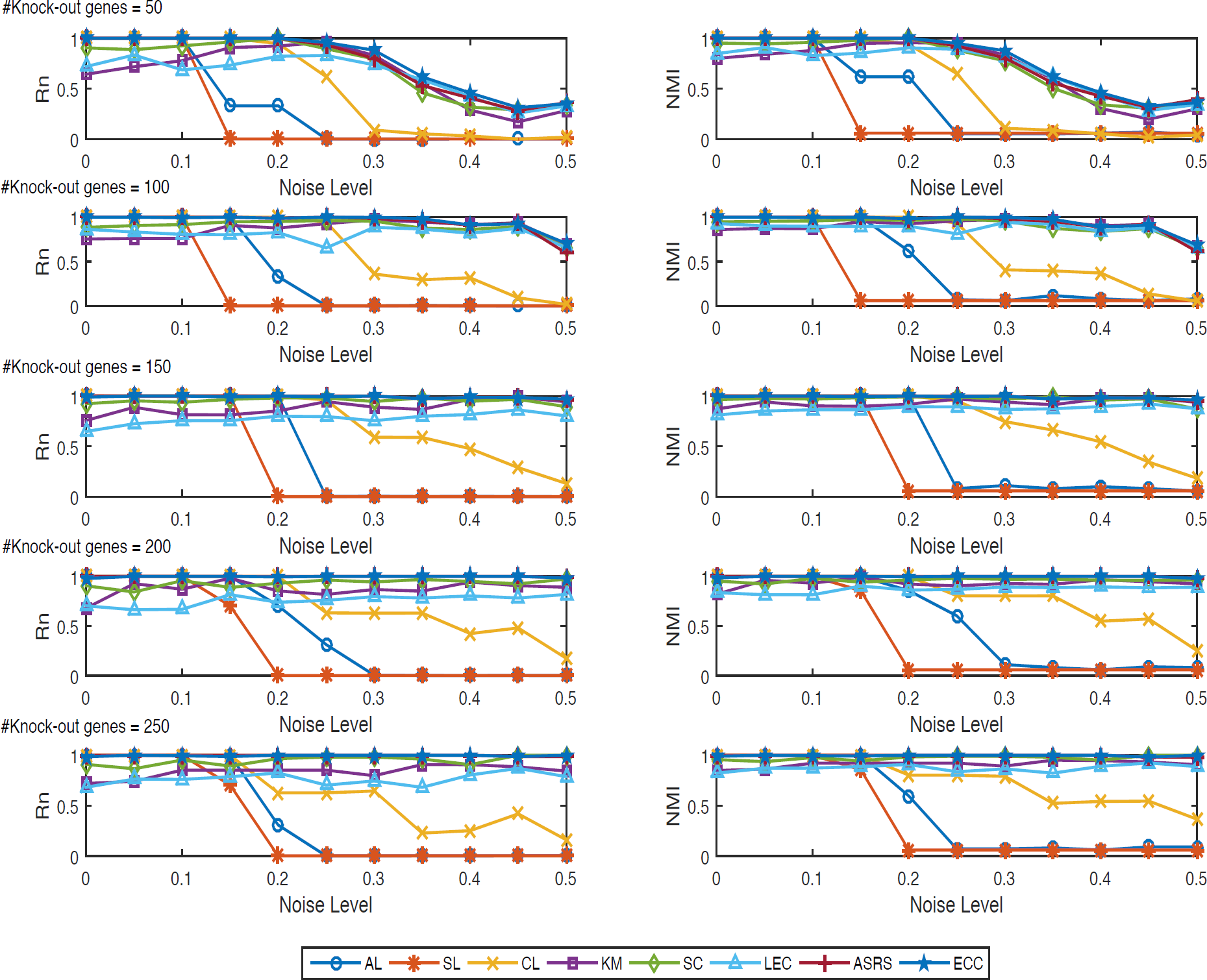
Performance of different clustering algorithms on the 55 synthetic datasets (based on an Erdös-Rėnyi random gene regulatory network of 500 genes). ECC has substantial advantages over other methods on the datasets marked by blue pentagram lines.

**Figure S4.**
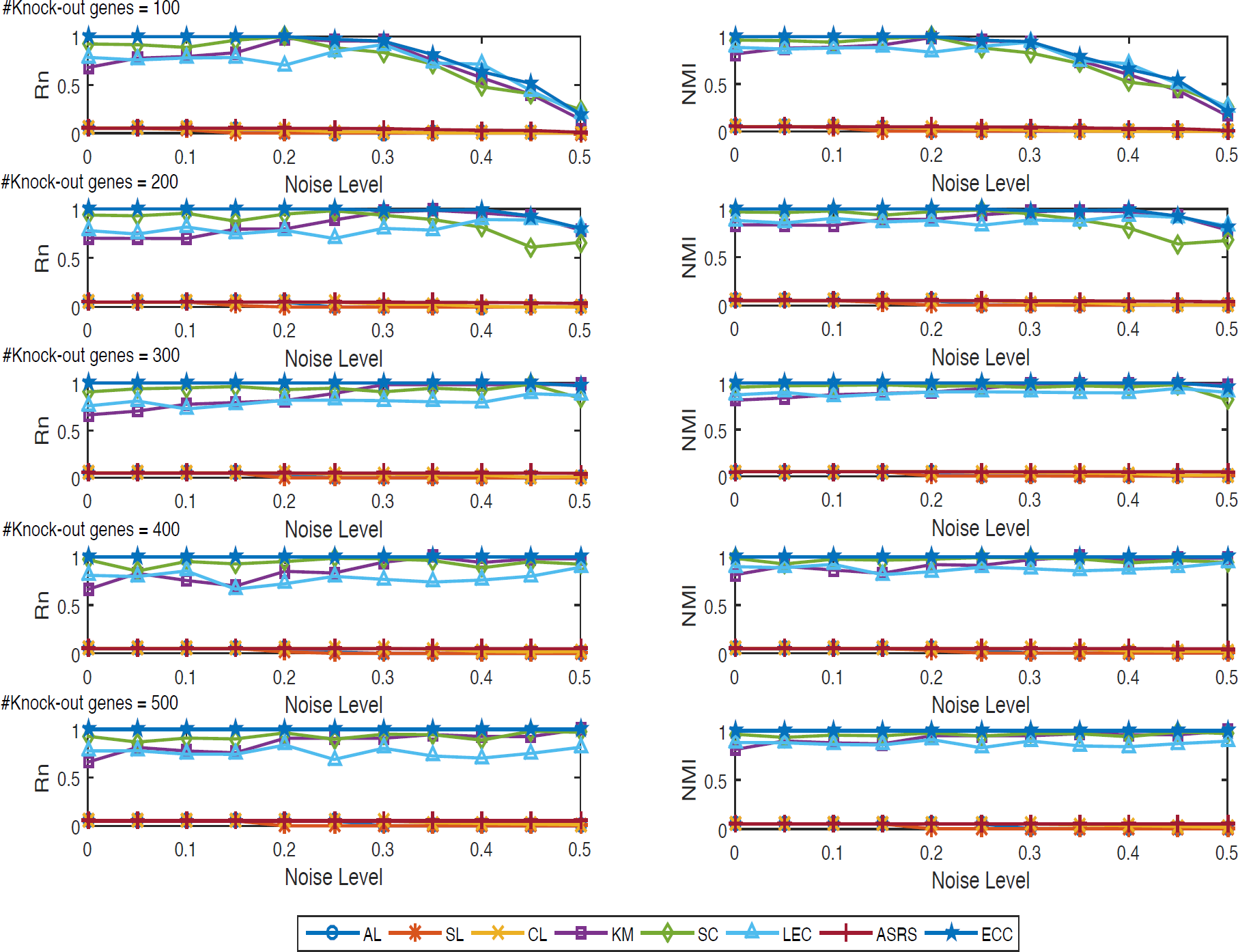
Performance of different clustering algorithms on the 55 synthetic datasets (based on a real human transcriptional regulation network of 2723 genes). ECC has substantial advantages over other methods on the datasets marked by blue pentagram lines.

**Figure S5.**
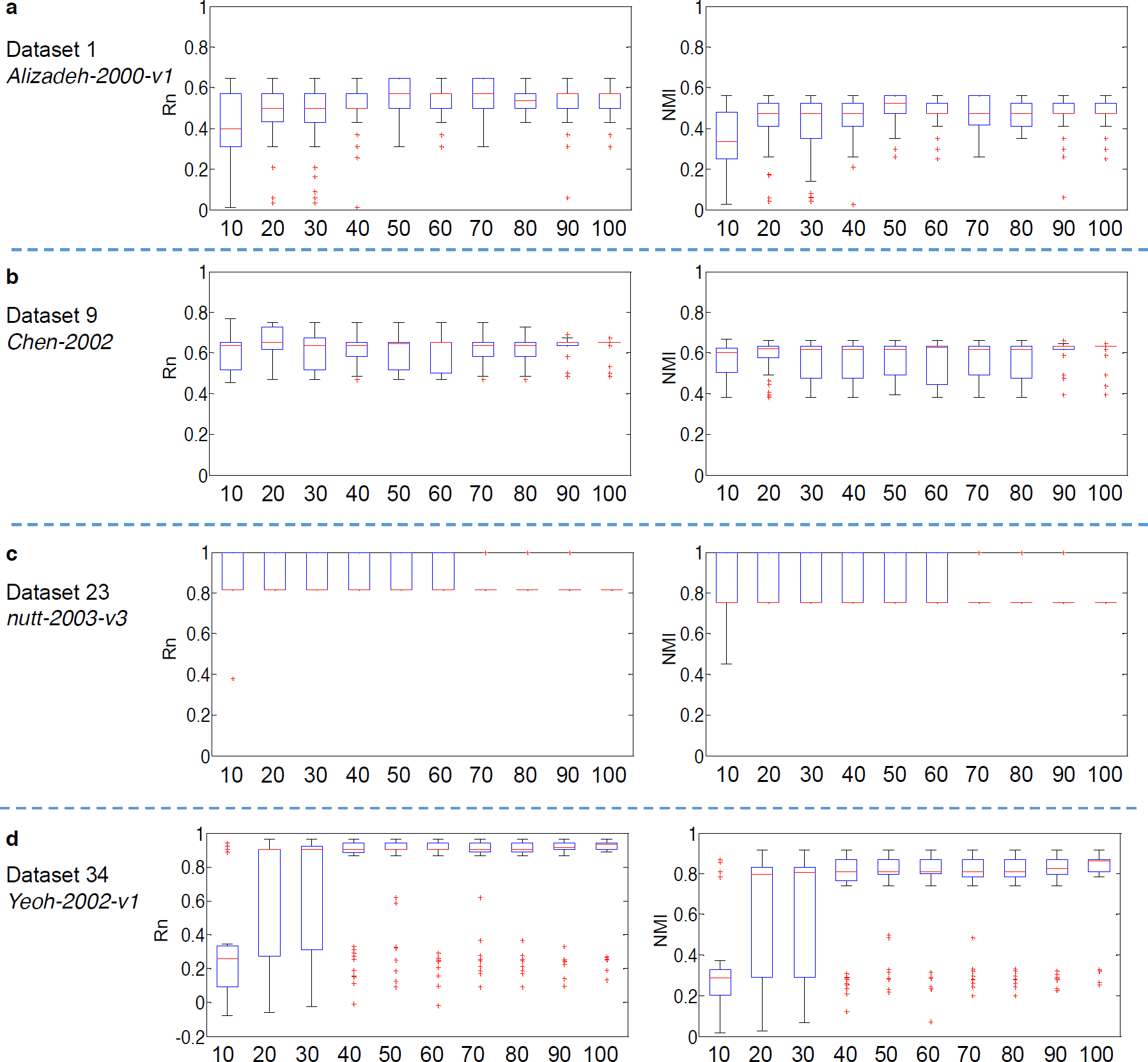
Impact of different numbers of basic partitions on ECC. The x-axis denotes the number of basic partitions. For each scenario, ECC runs 100 times for the boxplot. As the increase of basic partitions, the performance goes up and the variance becomes narrower and narrower.

**Figure S6.**
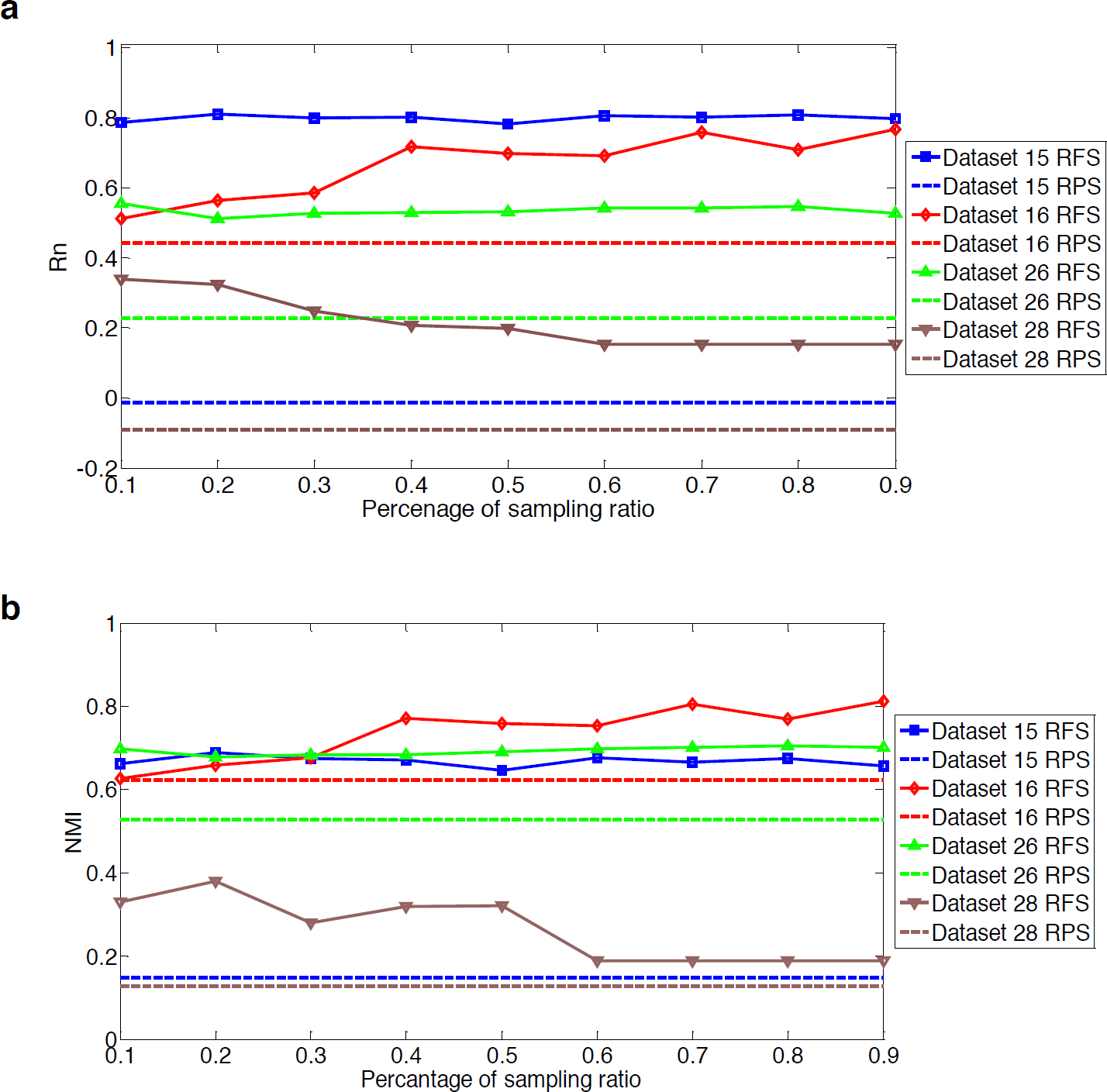
Random feature selection (RFS) strategies with different sampling ratios on ECC. On these four datasets, the performance of RFS exceeds those of RPS with all sampling ratios, which indicates that RFS can help to avoid noisy and irrelevant genes. Although it is difficult to select discriminative genes for cluster analysis, ECC can fuse the partial knowledge from RFS to achieve promising results.

**Figure S7.**
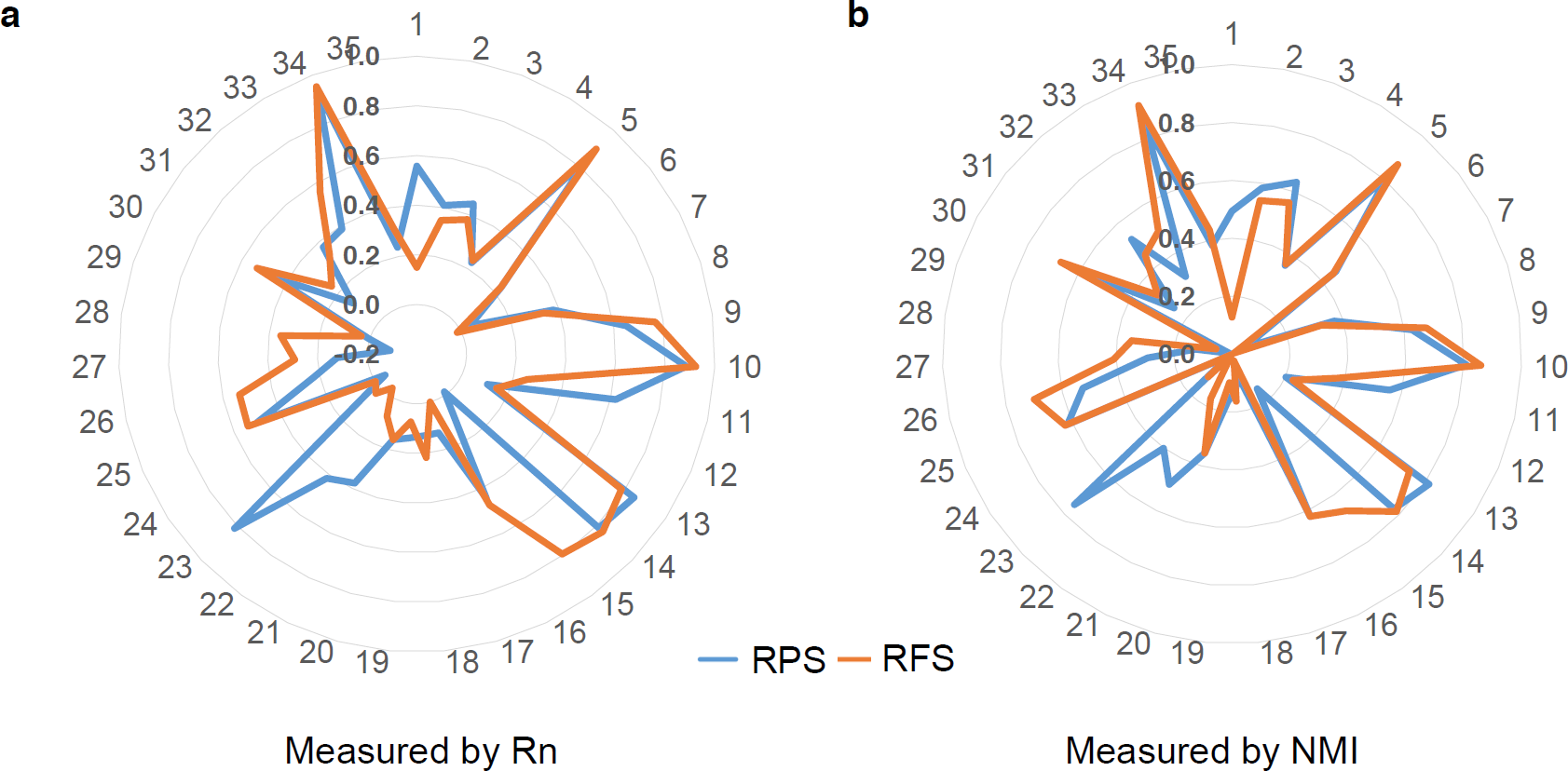
Comparison of different basic partition generation strategies on ECC. In RPS strategy, we apply *K*-means with the cluster number varying from 2 to 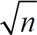. In RFS strategy, we apply *K*-means to 10% sampling ratio of the genes. RPS is suitable for the datasets which contain some potential sub-clusters, while RFS is suitable for the datasets containing noisy and irrelevant genes.

**Figure S8.**
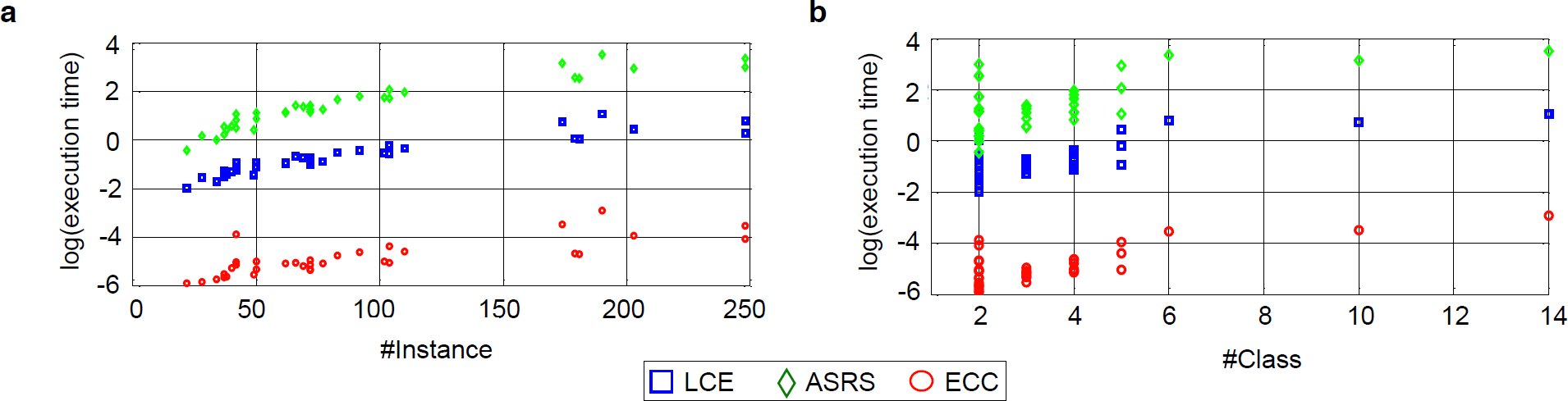
Execution time of three different consensus clustering methods. **a** shows the logarithm of the execution time in terms of the number of subjects and **b** shows the the logarithm of the execution time in terms of the cluster number. From these figures, ECC shows dramatically merits over two other consensus clustering methods (LCE and ASRS) in terms of efficiency. From the scope of these scatter plots, we can see that the time complexity of ECC is linear the number of subjects and the cluster number, while LCE and ASRS suffer from *O*(*n*^2^log*n*) and *O*(*n*^3^), respectively. This indicates ECC is suitable for large-scale gene expression data analysis.

**Figure S9.**
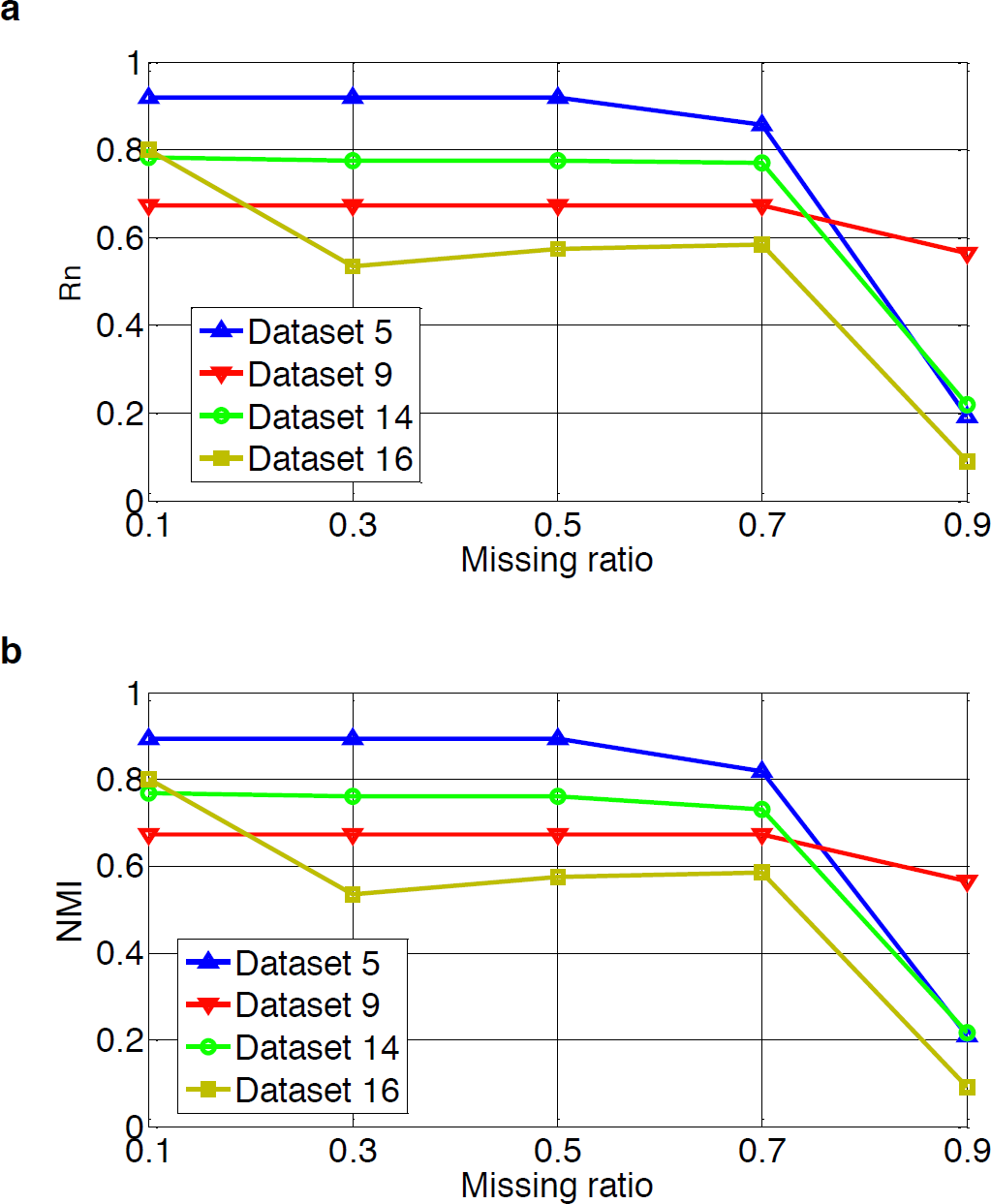
Performance of ECC with different missing ratios on 4 datesets. To generate incomplete basic partitions, we randomly remove some instances and employ K-means to assign the rest instances from 1 to *K*, where *K* is the user-defined cluster number. For these unsampled instances, the labels are assigned to be 0. The above process are repeated *r* = 100 times and we employ ECC to fuse these IBPs into a consensus one.

